# Inhibitory effect of *Bacillus subtilis* WL-2 and its IturinA lipopeptides against *Phytophthora infestans*

**DOI:** 10.1101/751131

**Authors:** Youyou Wang, Congying Zhang, Lufang Wu, Le Wang, Wenbin Gao, Jizhi Jiang, Yanqing Wu

**Affiliations:** College of Life Science, Hebei University, Baoding 071002, China; College of Biochemical and Environmental Engineering, Baoding University, Baoding 071000, China)

**Keywords:** Lipopeptides, IturinA, *Bacillus subtilis*, *Phytophthora infestans*, Inhibition

## Abstract

Potato late blight triggered by *Phytophthora infestans* ((Mont.) de Bary) represents a great food security threat worldwide and is difficult to control. Currently, *Bacillus* spp. have been considered biocontrol agents to control many fungal diseases. Here, *Bacillus subtilis* WL-2 was selected as the antifungal strain with the most potential against *P. infestans* mycelium growth. Additionally, the functional metabolites extracted from WL-2 were identified as IturinA-family cyclic lipopeptides (CLPs) via high-performance liquid chromatography (HPLC) and electrospray ionization mass spectrometry (ESI-MS). Analyses using scanning and transmission electron microscopy (SEM and TEM) revealed that IturinA caused a change in the mycelial surface and damage to the internal cell structure, including cell membrane disruption and irregular organelle formation. Moreover, propidium iodide staining and nucleic acid and protein release were detected to clarify the cell membrane damage caused by IturinA. Additionally, IturinA triggered reactive oxygen species (ROS) generation and malondialdehyde (MDA) production. Mitochondrial membrane potential (MMP), mitochondrial respiratory chain complexes activity (MRCCA), respiratory control rate (RCR), and oxidative phosphorylation efficiency (P/O) assays indicated that *P. infestans* mitochondria affected by IturinA were so seriously damaged that the MMP and MRCCA declined remarkably and that mitochondrial ATP production ability was weakened. Therefore, IturinA induces cell membrane damage, oxidative stress, and dysfunction of mitochondria, resulting in *P. infestans* hyphal cell death. As such, the results highlight that *B. subtilis* WL-2 and IturinA have great potential as candidates for inhibiting *P. infestans* mycelium growth and controlling potato late blight.

**IMPORTANCE:** Potato (*Solanum tuberosum* L.) is the fourth most common global food crop, and its planting area and yield increase yearly. Notably, in 2015, China initiated a potato staple food conversion strategy, and by 2020, approximately 50% of potatoes will be consumed as a staple food. The plant pathogen fungus *Phytophthora infestans* ((Mont.) de Bary) is the culprit of potato late blight; however, biological agents rather than chemicals are highly necessary to control this threatening disease. In this study, we discovered an antifungal substance, IturinA, a lipopeptide produced by *Bacillus subtilis* WL-2. Moreover, our research revealed the actual mechanism of IturinA against *P. infestans* mycelium growth and clarified the potential of *B. subtilis* WL-2 and IturinA as a biocontrol agent against *P. infestans* mycelium growth as well as for controlling the development of late blight in potato cultivation.

## INTRODUCTION

Behind rice, wheat, and corn, potato (*Solanum tuberosum* L.) is the fourth most stable food crop in the world. However, potato production is often endangered by many pathogens. Worryingly, late blight triggered by *Phytophthora infestans* ((Mont.) de Bary) could directly reduce or even eliminate potato production, and an outbreak of this disease could even result in a grievous economic loss in the agriculture industry(1, 2). At present, controlling late blight is achieved mainly using disease-resistant varieties and spraying chemical pesticides(3). However, due to the rapid variability of *P. infestans* and increase in physiological complexity of races, superphysiological races (R1-R11), which can overcome all the late blight protection genes (1.2.3.4.5.6.7.8.9.10.11), have emerged(4, 5). Additionally, as the result of the excessive use of chemicals, resistance of *P. infestans* to chemical pesticides has become increasingly common. In summary, the chemicals used have posed a massive challenge to control potato late blight and resulted in a great threat to food safety and the ecological environment(6). Therefore, exploration of suitable measures to control potato late blight is urgent. Surprisingly, biocontrol agents (BCAs), including microorganisms and secondary metabolites, have been further researched and even considered promising and environmentally friendly alternatives to the chemicals(7). With biocontrol method development, various antibiotic peptides, including polymyxin, daptomycin, chromobactomycin, subtilin and subtilosin, which were all extracted from *Bacillus* spp., have been considered potential future drugs based on their broad range of antibiotic activity, reduced toxicity, and safety to our environment(3, 7, 8).

With obvious biocontrol properties, cyclic lipopeptides (CLPs) synthesized from *Bacillus* spp. have been a focus of research in recent years. Additionally, CLPs with a wide range of antibacterial activities are one of the most abundant and highly yielded metabolites from *Bacillus* spp.(9). For its structure, CLPs consists of a peptide cycle composed of different amino acid arrangements and a lipid component composed of fatty acid chains of different lengths, and its molecular weight is approximately 1.1 kDa to 1.5 kDa(3). The structure of the peptide cycle combines with a long fatty acid chain to produce an amphiphilic trait, which determines the most suitable target sites on the cellular membranes(10). In addition, due to its variety and the number of amino acids as well as the diversity of fatty acid chain length, CLPs have multiple homologs(11). Moreover, CLPs derived from *Bacillus* spp. can be classified into three main subfamilies: iturin, surfactin, and fengycin(11–13). Interestingly, surfactin has powerful antiviral activities but low activities against bacteria and fungi(14), while iturin and fengycin exhibit an obvious antifungal activity against a range of filamentous fungi(9, 15). Most of the special biocontrol mechanisms of iturin and fengycin against phytopathogens have been characterized(16). Specifically, fengycin derived from *Bacillus subtilis* BS155 has a strong antagonistic activity against *Magnaporthe grisea* by reactive oxygen species (ROS), chromatin condensation, and separation of cell walls from the membranes(17). Many results have shown that iturin can inhibit the mycelium growth of many fungi, including *Candida albicans*, *Aspergillus flavus*, *Sclerotinia sclerotiorum*, *Botrytis cinerea*, *Monilinia fructicola*, *Fusarium graminearum* and so on(18–21). More specifically, iturin extracted from *Bacillus amyloliquefaciens* FZB42 could cause morphological changes in the plasma membranes and cell walls of *F. graminearum* hyphae, lead to ROS accumulation, and induce cell death in conidia (22). When iturin was used against *S. sclerotiorum*, it could also trigger a separation of cell walls from membranes and even form a pore in the cell membrane, resulting in leakage of the cytoplasm(19). Additionally, IturinA disrupted the *B. cinerea* cytoplasmic membrane; created transmembrane channels, resulting in K^+^ leakage; prevented spore germination; and impaired mycelium development(23). CLPs produced by *Bacillus* have antiphytopathogen activities; however, the inhibitory effect of lipopeptides on *P. infestans* remains poorly understood(3). Therefore, through this study, we intended to compare the potential of three bacteria, *B. subtilis* WL-2 (MK241790), *Pseudomonas fluorescens* WL-1 (MH229994) and *Bacillus pumilus* W-7 (KX056277), as efficient BCAs for the control of *P. infestans* mycelium growth. Meanwhile, CLPs extracted from the bacteria were purified using high-performance liquid chromatography (HPLC) and identified using Fourier transform infrared spectroscopy (FTIR) and mass spectrometry (MS). In addition, the antifungal mechanism of CLPs on *P. infestans* mycelium growth, cell integrity, mitochondrial damage, and ROS generation were exploited to evaluate the consideration of CLPs as antifungal agents against potato late blight in the future.

## 1 MATERIALS AND METHODS

### 1.1 Inhibition effect of three strains against *P. infestans*

The fungal pathogen *Phytophthora infestans* ((Mont.) de Bary) W101 was obtained from the China General Microbiological Culture Collection Center (CGMCC) and grown on rye (R) solid medium at 20°C in the dark(24). *Bacillus subtilis* WL-2 (MK241790), *Pseudomonas fluorescens* WL-1 (MH229994) and *Bacillus pumilus* W-7 (KX056277) bacteria were isolated from *Capsicum frutescens* leaves and cultured on Luria Bertani (LB) solid medium at 35°C(25). Living cell (LC) of bacteria was grown on LB solid medium and incubated for 24 h at 37°C. To obtain cell suspension (CS), LB liquid medium was inoculated with each strain and incubated for 20 h at 37°C and 200 rpm, and the final concentration (1×10 ^7^ CFU/mL) was regulated by distilled water. Prepared 2% seed culture (SC, 1×10 ^7^ CFU/mL) were transformed into flasks containing 100 mL of LB liquid medium and incubated at 30°C and 200 rpm for 96 h. Finally, the liquid culture was centrifuged (10,000 × g, 4°C) for 10 min, and LC were filtered using cellulose acetate membranes (0.22 µm) to obtain a cell-free supernatant (CFS) solution (26, 27). Eight-day-old *P. infestans* mycelium was scraped into 10 mL of distilled water and oscillated to expose sporangium; then, the sporangium suspension was regulated to 1×10 ^7^ CFU/mL using distilled water. Finally, the sporangium suspension was released at 10°C for 3 h to obtain zoospores, and the zoospore solution was regulated by sterile water up to 1×10 ^7^ CFU/mL for further analysis(28).

The inhibitory effect of the three strains on the growth of *P. infestans* mycelium was assessed on LC, CS, and CFS using the plate dual culture method(29). First, a *P. infestans* mycelium disk (diameter = 7 mm) was placed on the center of R solid medium (diameter = 9 cm) and cultivated for three days in advance. Then, LC was placed at a position 3 cm away from the center disk, and an equal volume of blank LB liquid medium was placed as the control. Additionally, the punch method(24) was adopted to determine the inhibitory effect of CS and CFS. Similarly, every punch (9 mm) was added with 100 µL of CS and CFS, and an equal volume of blank LB liquid medium was added as the control. Finally, after coincubation at 20°C for five days, the inhibitory zones (mm) were measured using the cross method(24), and the inhibition rate was determined by the described formula:

> Inhibition rate (%) = (C − T) / C× 100(30)

Where C represents the fungal colony radius of the control, and T symbolizes the radius of the treatment group.

### 1.2 Biocontrol assays for the WL-2 strain

A potato-sensitive variety of “Bintje” was used to prepare *in vitro* tuber slices (2.0 cm×2.0 cm×0.5 cm) and healthy leaves(31). Treatment measures, such as disease prevention (DP), simultaneous inoculation (SI), and disease therapy (DT), were conducted to evaluate the biocontrol effect of the WL-2 strain. For DP, a CS (1×10 ^6^ CFU/mL, 50 µL per slice) was smeared over the potato tuber slices and leaves at room temperature in advance. Then, an equal volume of blank LB liquid medium was smeared as the control. After 48 h, a *P. infestans* mycelium disk (diameter =7 mm) was placed on the top of the tubers, and the *P. infestans* zoospore suspension (1 x 10^7^ CFU/mL, 20 µL per slice) was added at the back of the leaves. Finally, tuber slices were cultured for six days after inoculation at 20°C in the dark, and the leaves were cultured for six days after inoculation at 20°C in a 16 h light/8 h dark photocycle. For SI, the CS and *P. infestans* were inoculated at the same time. For DT, *P. infestans* was inoculated two days in advance, and then the CS was processed(32). Based on a 1-9 scale, the disease index was calculated according to the following formula:

> Disease index = ∑(d_i_×l_i_)/(L×N)×100(30, 33)

Where d_i_ represents the disease grade, and the number of leaves or tubers at different grades were represented with l_i_. L symbolizes the sample number, and N indicates the highest disease grade.

### 1.3 Detection of CLPs production ability and preparation of crude lipopeptide extract (CLE)

**(1): Identification of hemolysis ability** To determine the hemolysis characteristics of CLPs, the WL-2 strain was inoculated on sheep blood medium at 37°C for 48 h to detect hemolysis activity, and inoculation of *Escherichia coli* (without hemolysis ability) was used as a control(34). **(2): CFS surface tension (ST).** CFS was detected every 12 h for 96 h (8 times), and ST was recorded with a tensiometer (Gibertini, Milan, Italy) using the Wilhelmy plate method(35). The instrument was calibrated against distilled water (ST = 73.1 mN/m) for accurate measurements(36). **(3): Oil dispersal diameter measurement**(37). Soybean oil (1 mL) was added above the surface of distilled water (30 mL, 4°C), and then a white oil film on the distilled water surface was formed. WL-2 CFS (50 μL) was added to the center of the oil film to record the diameter of the oil dispersal ring. Meanwhile, blank culture solution was used as a control. **(4): Emulsification index determination**(38). CFS (3 mL) and soybean oil (3 mL) were mixed in a tube, and then the mixture was treated with an ultrasonic cleaning instrument (SK2510, KUDOS, Shanghai, China) for 1 min to mix thoroughly. Finally, the mixture was incubated statically for 24 h, the height of the emulsion layer was measured, and the percentage of emulsifying properties was calculated as follows:

> Emulsifying properties (%) =(emulsion layer height/total height)×100(39)

The WL-2 strain inoculated into LB liquid medium was cultured at 35°C and 180 rpm for 24 h to prepare an SC (1×10 ^6^ CFU/mL). Then, 3% (by vol) SC was transferred into the flask (1,000 mL) containing 400 mL of Landy liquid medium(40) and cultured at 30°C and 180 rpm for 96 h. The acid precipitation method(41) was used to prepare CLE. The culture was centrifuged (10,000 × g, 4 °C) for 10 min to remove LC. Then, 6 N HCL was added to adjust the pH of the supernatant (pH=2.0) and induce precipitation(42). Finally, the CLPs contained in the precipitation were fully dissolved in methanol, and a rotary evaporator (RE52CS-1, YARONG, Shanghai, China) was then used at 50°C and 65 rpm to obtain CLE for further analysis.

### 1.4 MALDI-TOF-MS and antifungal assays

The CLE methanol solution (1 mg/L, 10 µL) was mixed with 1 µL of saturating matrix solution of α-cyano-4-hydroxy-cinnamic acid. The matrix solution containing TFA (0.1%) was prepared using H_2_O and CH_3_CN (1:1, v/v). Based on a 20 kV accelerating voltage, the samples were detected, and matrix-assisted laser desorption ionization time-of-flight mass spectrometry (MALDI-TOF-MS, AUTOFLEX III, Bruker Daltonics) was utilized to analyze the sample in positive mode. Finally, the *m/z* values in the range from 600 to 1,700 were analyzed (16, 43–45).

The disk diffusion(43) method was adopted to evaluate the antioomycete activity of CLE. A *P. infestans* disk (7 mm) was incubated on R solid medium plates for three days in advance, and filter paper disks (5 mm) containing 6 µL of CLE solution (1, 3, and 5 mg/mL) were then placed at a position 4 cm away from the center disk. Meanwhile, the same volume of the fungicide Metalaxyl (15 µg/mL) and a methanol solution were used as the control. The plates were co-incubated at 20°C for five days, and inhibition rates were determined(30).

### 1.5 HPLC and FTIR analysis

Commercial standard lipopeptides (surfactin and iturin, Sigma-Aldrich, United States) and CLE methanol solution (10 mg/L) were run on an HPLC system (Waters, E2695, United States) with a C_18_ reverse-phase column (5 µm, 4.6 × 150 mm) under the same conditions(46). Water and acetonitrile were selected as the mobile phase at a ratio of 20:80 by volume. The injection volume was 1 mL per min, and the eluate was monitored at 214 nm. According to the peak times of standard lipopeptides, the peaks of potential lipopeptides contained in CLE were collected and dried at room temperature(45, 46).

According to the matched peaks of CLE, the functional groups presented in the CLE were determined using FTIR(47). First, 1 mg of every purified lipopeptide (matched peaks) and standard lipopeptide was ground in KBr (100 mg, spectral grade) to prepare translucent pellets(48). Data from the FTIR spectrum were collected between 500 and 4,000 cm^-1^, and the characteristic absorbance peaks were analyzed(49).

### 1.6 MALDI-TOF-MS/MS and antifungal activity analysis

As described in the MALDI-TOF-MS method, 1 µg/mL purified lipopeptide methanol solutions (peak a and peak b) were detected using MALDI-TOF-MS/MS (MALDI-TOF, AUTOFLEX III, Bruker Daltonics) coupled with HCD mode to clarify the amino acid sequence in lipopeptides(50). Depending on the precursor ion of interest, a suitable collision energy was used from the range of 35 to 50 eV(51). Antioomycete activity of purified surfactin and IturinA was evaluated using the disk diffusion method(43). Similarly, 6 µL of purified surfactin and IturinA solution (dissolved in distilled water) with different concentrations (20, 30, 40 and 50 µg/mL) was added to filter paper disks (5 mm), and distilled water was used as a control. After coincubation at 20°C for five days, the inhibition zone and inhibition rate were determined.

### 1.7 Inhibition effect of IturinA against *P. infestans*

#### (1) The recovery of *P. infestans* mycelium and sporangium after inhibition

*P. infestans* marginal mycelium disks (diameter = 7 mm) inhibited by IturinA (20, 30, 40, and 50 µg/mL) were transferred onto fresh R solid medium, and mycelium disks without inhibition were used as a control. All the treated plates were incubated at 20°C for seven days in the dark, and the colony diameter and growth rate were calculated according to the formula below.

Recovery growth rate (%) = (The maximum colony diameter / Total days) × 100 (24) Meanwhile, the sporangia inhibited by IturinA (20, 30, 40, and 50 µg/mL) were separated using screen mesh (diameter = 50 µm) and adjusted to 1×10 ^7^ CFU/mL using distilled water. Finally, the sporangium suspension was induced to release zoospores at 4°C for 3 h in the dark, and sporangium direct germination was induced at 25°C for 5 h in the dark(52). An optical microscope (OM, BX53, OLYMPUS, Japan) was used to observe 300 spores to calculate the zoospore release rate and sporangium direct germination rate according to the formula below:

> Release or germination rate (%) = (Total release or germination number /Number of total spores) × 100(52)

#### (2) Optical microscopy (OM), Scanning Electron Microscopy (SEM) and Transmission Electron Microscopy (TEM) observation

The marginal mycelia of IturinA (50 µg/mL) -inhibited *P. infestans* were collected and washed twice in PBS (pH 7.2), and then an OM system was utilized to observe mycelium damage via morphology(24). Meanwhile, mycelia were fixed using 2.5% (v/v) glutaraldehyde (Solarbio, Beijing, China) for 24 h and dehydrated for 30 min in every step using aqueous ethanol solutions (30, 50, 70, and 90%, v/v). Then, morphological and surface changes were observed using an SEM system (JSM-7500F, JEOL, JAPAN)(53). A TEM (JEM-2100F, JEOL, JAPAN) system(53, 54) was also adopted to evaluate the structural characteristics of inhibited mycelia. Similar to above, 2.5% (v/v) glutaraldehyde was used to fix damaged mycelia, 1% (v/v) osmium tetroxide was used to fix mycelia at 20°C for 20 min, and finally, a microtome (YD335, Leica, Germany) was used to prepare thick specimens (70 nm) for TEM observation.

#### (3) *P. infestans* cell membrane damage induced by IturinA

*P. infestans* marginal mycelia and sporangia inhibited by IturinA (50 µg/mL) were collected and washed twice with 20 mM PBS buffer (pH 7.2). Then, 30 µM propidium iodide was used to stain cells in an ice bath for 10 min. Additionally, a group without inhibition was used as a control(55). Subsequently, mycelia were observed using a filter (535 nm/615 nm) under a confocal fluorescence microscope (CFM, FV3000, OLYMPUS, Japan)(56).

Changes in membrane permeability caused by IturinA were investigated in a mycelium-soaked solution according to the changes in electrical conductivity and optical density at 260 nm and 280 nm. First, *P. infestans* marginal mycelia (100 mg) inhibited by IturinA (50 µg/mL) were collected into a plate and then washed twice with distilled water (20 mL). Filter paper was used to remove water drops mixed with mycelia, and then, the prepared mycelium sample was resuspended in 10 mL of distilled water. Mycelia without inhibition were treated as a control. Cell membrane permeability was determined using a conductivity meter (S7-Meter, METTLER TOLEDO, Switzerland) according to the electrical conductivity of the mycelium solution after being suspended for 0, 20, 40, 60, 80 and 100 min, respectively. A mycelium solution boiled for 10 min was considered a control group (final conductivity). Finally, the relative conductivity of the mycelium was calculated according to the following formula:

> Relative conductivity (%) = (Conductivity/Final conductivity) × 100(57)

Additionally, the absorbances of the mycelium solution at 260 and 280 nm were measured by an ultraviolet-visible light detector (UV-1800, SHIMADZU, Japan)(58, 59) to assess nucleic acid and protein leakage. The measurement was conducted at regular intervals of 20 min, from 0 min to 100 min (6 times), and the significant difference was compared with that of the control group(60).

### 1.8 IturinA leads to the accumulation of Reactive Oxygen Species (ROS) and Malondialdehyde (MDA) production

ROS accumulation in *P. infestans* cells induced by IturinA was detected with DCFH-DA, which is commonly used to evaluate oxidative stress in cells(61). First, five *P. infestans* mycelium disks (diameter =7 mm) were transferred into a flask containing 100 mL of R liquid medium and cultured at 20°C and 180 rpm for 48 h. Afterward, IturinA (50 µg /mL, final concentration) was added to the mycelium suspension and incubated for 0, 4, 8, 12, 16, 20, and 24 h. Additionally, the ROS-inducing drug Rosup (10 µg/mL, final concentration) was used to treat for 20 min and considered a positive control, while distilled water was used as a negative control. Subsequently, *P. infestans* mycelium was resuspended in PBS buffer (pH 7.2), and then 10 μM DCFH-DA was co-incubated with mycelia for 20 min. Finally, the CFM system was used to analyze fluorescence intensity(62).

MDA is the most important product marker of ROS, so detection of the MDA concentration was performed to assay ROS intensity and cell damage. After treatment with IturinA, mycelium was analyzed using MDA assay kits (Beyotime Biotechnology, China), and the absorbance at 532 nm was measured to assay MDA production using an ultraviolet-visible light detector (UV-1800, SHIMADZU, Japan)(60).

### 1.9 IturinA leads to mitochondrial damage

#### (1) Assay of Mitochondrial Membrane Potential (MMP)

For determination of MMP (mtΔψ), a mitochondrion-specific lipophilic cationic fluorescence dye, JC-1, was used to assay MMP in *P. infestans* mycelium(63). Based on the results above, the ROS generation induced by IturinA reached the highest value when the incubation time was 16 h, so mycelium incubated for 16 h was collected and stained with 10 μg/mL JC-1 in the dark for 20 min. Next, the JC-1 solution was removed(62), and the mycelium was resuspended in PBS. The fluorescence of JC-1 (red fluorescence and green fluorescence) was monitored at Ex/Em = 490/525 nm and 490/590 nm using a CFM system(64).

#### (2) Effect of IturinA on Mitochondrial Respiratory Chain Complexes Activity (MRCCA), Respiratory Control Rate (RCR) and Oxidative Phosphorylation Efficiency (P/O)

After inhibition by IturinA (50 µg/mL) for 16 h, *P. infestans* mycelia were collected. Then, 5 mL of lysis buffer was added to suspend mycelia and extract mitochondria according to the Mitochondrial Isolation Kit (Beyotime, Shanghai, China) instructions (65). Next, mitochondrial oxidative phosphorylation detection was conducted after mitochondrial disruption through four freezing (-80°C) and thawing (30°C) cycles(66, 67). The MRCCA, including that of complex I-V, was measured based on the absorbance decline at different values(68, 69). The oxidation rate of NADH catalyzed by complex I was evaluated according to the absorbance decline at 340 nm to reflect complex I activity. In the complex II-catalyzed succinic acid oxidation reaction, DCPIP (2,6-dichlorophenol indophenol) was used as a coloring agent, and the reduction in absorbance at 600 nm was considered the activity decline of complex II. Complex III activity was detected according to the reduction rate of ferricytochrome c by CoQ_2_ (absorbance at 550 nm), and complex IV activity was evaluated as the cyanide-sensitive oxidation of ferrocytochrome c (absorbance at 550 nm). The activity of complex V was reflected by measuring the oxidation rate of NADH (absorbance at 340 nm). Mitochondrial respiratory chain complexes I-V enzyme activity was detected according to the kit instructions (AmyJet Scientific, Wuhan, China). However, mycelia without inhibition were treated as a control.

One milligram of inhibited mycelium (IturinA treated for 16 h) was placed in a respirator (O2k-FluoRespirometer, Oroboros, Austria) pool containing 2 mL of respiratory solution. Then, 2 mol/L glutamic acid (10 μL), 0.4 mol/L malic acid (5 μL), and 2.5 mmol/L succinic acid (100 μL) were added into the reaction pool, and subsequently, 2 μL of 100 mmol/L adenosine diphosphate was added to obtain STATE 3 respiration. At the time of ADP depletion, the respiratory rate was considered STATE 4; meanwhile, 1 μL of 1 mmol/L rotenone and 5 mmol/L (1 μL) antimycin A were added to inhibit respiration. Finally, the ratio of STATE 3 to STATE 4 was considered the RCR(65, 70). The ratio of ATP production to oxygen in the presence of respiring substrates and ADP was considered the P/O (71, 72). The treatment of mycelium without inhibition was assayed as a control.

All the different treatments above were repeated three times, and the final results are shown via the average value.

## 2 RESULTS

### 2.1 Comparison of the inhibition of *P. infestans* by three strains

The inhibitory effect of three strains against *P. infestans* is presented in Fig. 1 and Tab. 1. The LC of the three strains expressed a strong inhibitory effect on the growth of *P. infestans* mycelium, and all inhibition rates were above 60% (Fig. 1A, Tab. 1). In addition, the WL-2 strain had the strongest inhibitory effect (Fig. 1A-a), and the inhibition rate reached a maximum of 75.6%, which was significantly different from that of the other strains (*P*<0.05). Suppression of the growth of *P. infestans* mycelium by CS was stronger than that by LC treatment for all three strains (Fig. 1A, Fig. 1B, Tab-1), and the inhibition rates were all above 80%. Meanwhile, the inhibitory effect of the WL-2 CS was the most prominent, and the inhibition rate reached a maximum of 93.7%. After inhibition by the WL-2 CS, *P. infestans* mycelium could barely grow. Additionally, in the CFS experiment, the inhibition effect (inhibition rate was 80.7%) of the WL-2 strain was significantly better than that of WL-1 and W-7 (*P*<0.05). Altogether, the inhibitory effect of the WL-2 strain on the growth of *P. infestans* mycelium was significantly better than that of the other strains.

**Fig. 1.**
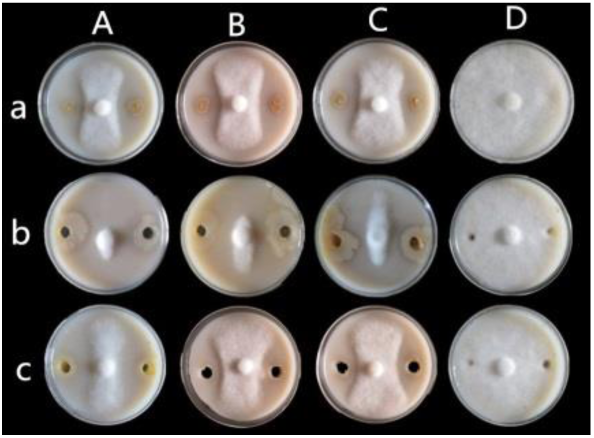
Comparison of three strains against *P. infestans*. Note: a: LC; b: CS; c: CFS; A: WL-2; B: WL-1; C: W-7; D: Control.

**Tab. 1.** Comparison of the inhibition of *P. infestans* by three *Bacillus* species.

| Strains | Inhibition rate (%) |  |  |
| --- | --- | --- | --- |
|  | LC | CS | CFS |
| WL-2 | 75.6 a | 93.7 a | 80.7 a |
| WL-1 | 65.4 b | 86.1 b | 56.6 b |
| W-7 | 62.5 b | 84.2 b | 58.7 b |
| CK | 0 c | 0 c | 0 c |

Note: The different letters a, b, c, and d in the same column symbolize a significant difference ( $P<0.05$ ), and the same is true below.

### 2.2 Biocontrol effect of WL-2 CS on tubers and leaves *in vitro*

With the most prominent inhibition effect against *P. infestans* mycelium growth, the WL-2 CS was selected to test the biocontrol effect on *in vitro* potato tissues. After six days of treatment using the CS alone, tubers (Fig. 2A-a) and leaves (Fig. 2B-a) were bright and without evident discoloration and decay, which indicated that the CS had no side effect on potato tissues. After DP, SI and DT treatments on tubers (Fig. 2A-b, c, d), the *in vitro* disease indices were 6.5, 16.2, and 35.4, respectively, which were significantly lower than those of the control (77.6, *P*<0.05, Fig. 2A-e). On leaves (Fig. 2B), the *in vitro* disease indices of DP, SI and DT (Fig. 2B-b, c, d) were 4.3, 10.9, and 25.3, respectively, which were also significantly lower than those of the control group (52.3, *P*<0.05). In addition, DP treatment was the best way to control late blight, and the disease index was the lowest compared with that of the SI and DT groups.

**Fig. 2.**
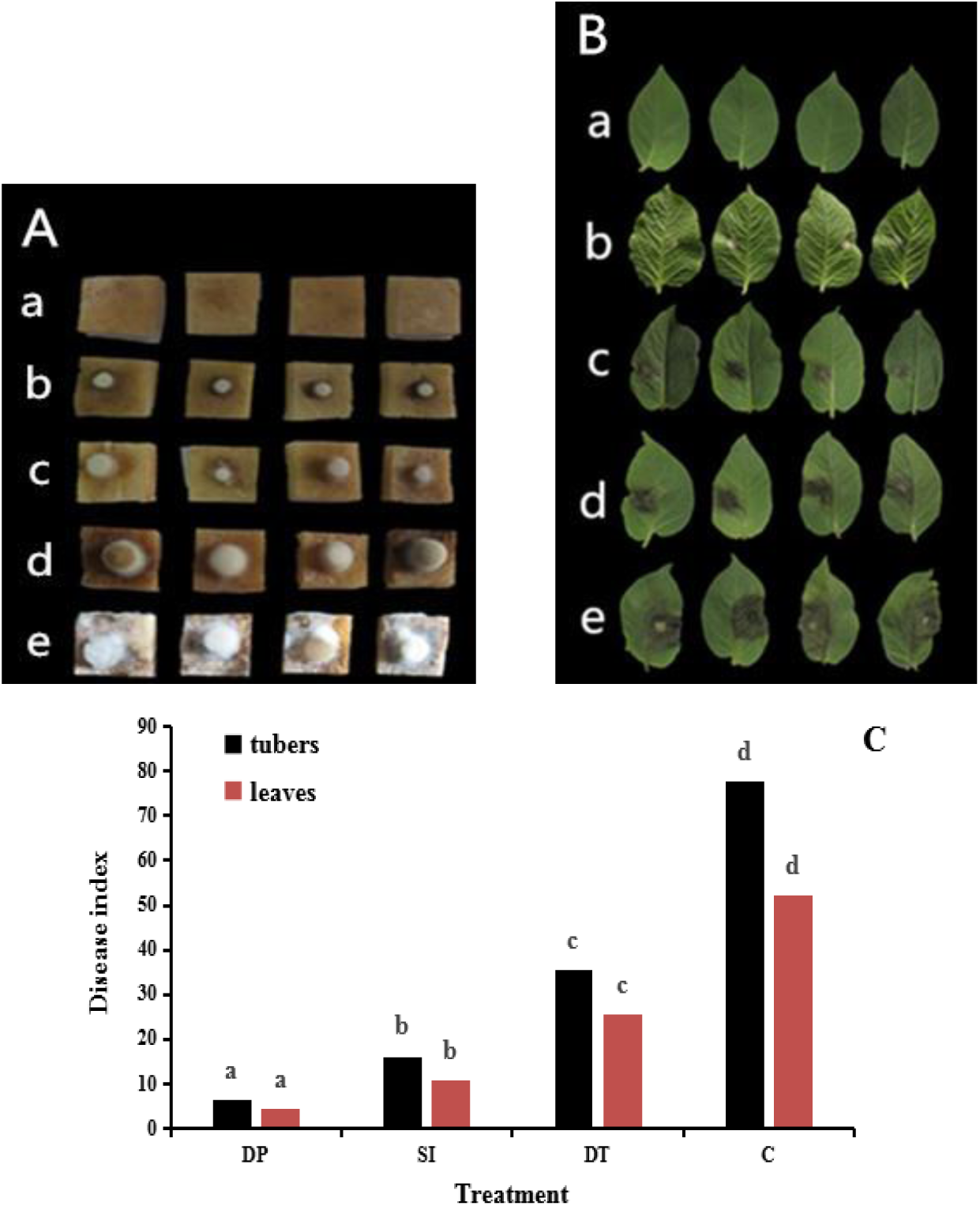
Biocontrol effect of the WL-2 CS on potato tissues *in vitro*. Note: A: *In vitro* tubers, A-a: Negative control (LB liquid medium); A-b: DP (disease prevention); A-c: SI (simultaneous inoculation); A-d: DT (disease therapy); A-e: Control (C, sterilized water); B: *In vitro* leaves; C: Comparison of disease indices. The different lowercase letters between different groups indicate a significant difference (*P*<0.05); the same is true below.

### 2.3 Detection of CLP production ability

**(1) Hemolysis activity:** The WL-2 strain inoculated on a sheep blood plate produced a transparent trace around the strain colony (Fig. 3A-a), while in the control group (*E. coli*), there was no transparent trace (Fig. 3B-a). The transparent trace indicated that the WL-2 strain had an obvious hemolysis activity. **(2) Detection of oil dispersal effect:** In the treatment group (WL-2 CFS), the oil film produced a large oil dispersal ring (diameter = 4.92 cm) in the plate center (Fig. 3A-b), while the oil dispersal ring that occurred in the control group was very small (diameter = 0.85 cm, Fig. 3B-b), and there was a significant difference between the two groups (*P*<0.05). **(3) Emulsification index:** The emulsification percentage was as high as 82.1% (Fig. 3A-c) in the treatment group, while the emulsification percentage of the control group was only 18.6% (Fig. 3B-c), and there was a significant difference (*P*<0.05) between the two groups. **(4) Measurement of ST:** ST properties are critical to the function of CLPs(73). With increasing WL-2 incubation time, the ST value was significantly reduced, and when the incubation time was 60 h, the ST value decreased from 73.1 mN/m (control) to the lowest value, 38.7 mN/m (Fig. 4).

**Fig. 3.**
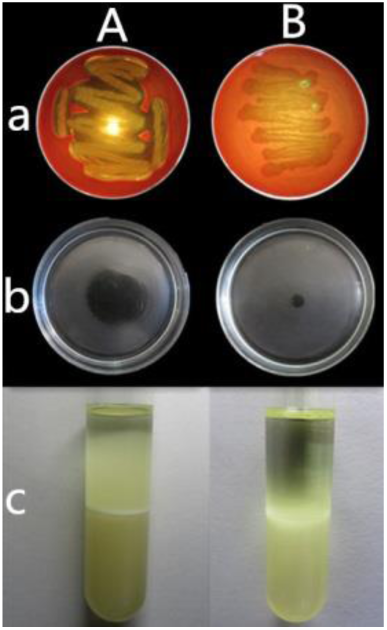
Determination of hemolysis, oil dispersal, and emulsification activities. Note: A: Treatment group; B: Control group. a: Hemolysis activity (*E. coli* as control); b: Oil dispersal diameter; c: Determination of emulsification index.

**Fig. 4.**
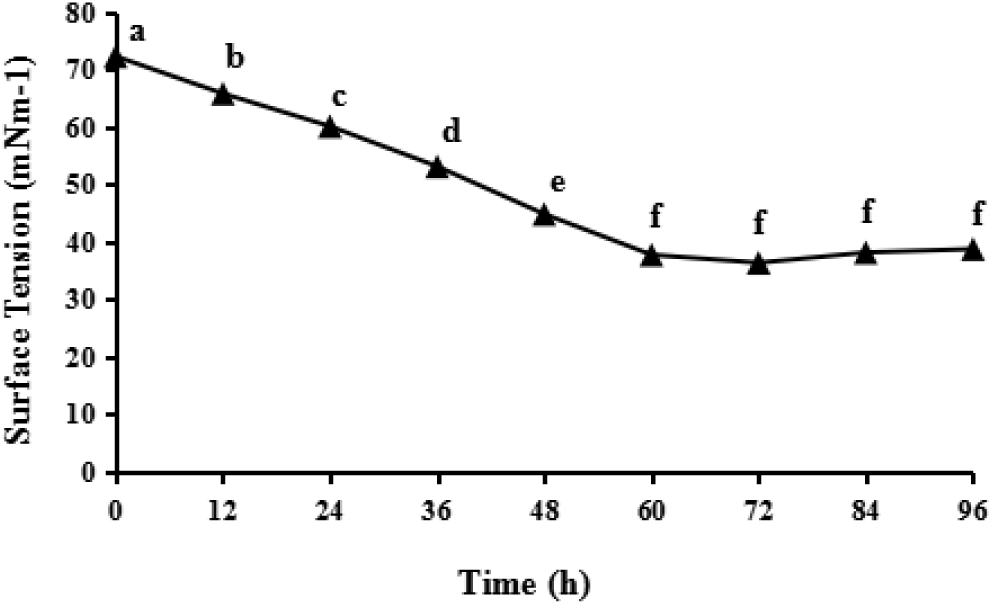
Measurement of ST. Note: The ST of distilled water was 73.1 mN/m.

### 2.4 MALDI-TOF-MS and antifungal assays

The average yield of prepared CLE was 2.3 g/L. Lipopeptides contained in the CLE appeared at molecular weights ranging from 1,000 to 1,100 (Fig. 5). The obvious molecular weights of 1,022.68, 1,036.69, and 1,050.71 were inferred to be surfactin (C_14_ - C_16_) with H+ adduct ions. In addition, the peaks at 1,044.66, 1,058.67, 1,072.69, and 1,086.70 were speculated to be surfactin (C_14_ - C_17_) with Na+ adduct ions (Fig. 5, Tab. 2). The molecular weights of 1,065.53 and 1,079.55 were considered to be IturinA with a fatty acid chain from C_14_ to C_15_ and with Na+ adduct ions (Fig. 5, Tab. 2). Based on the results above, lipopeptides of surfactin and IturinA contained in CLE were preliminarily determined.

**Fig. 5.**
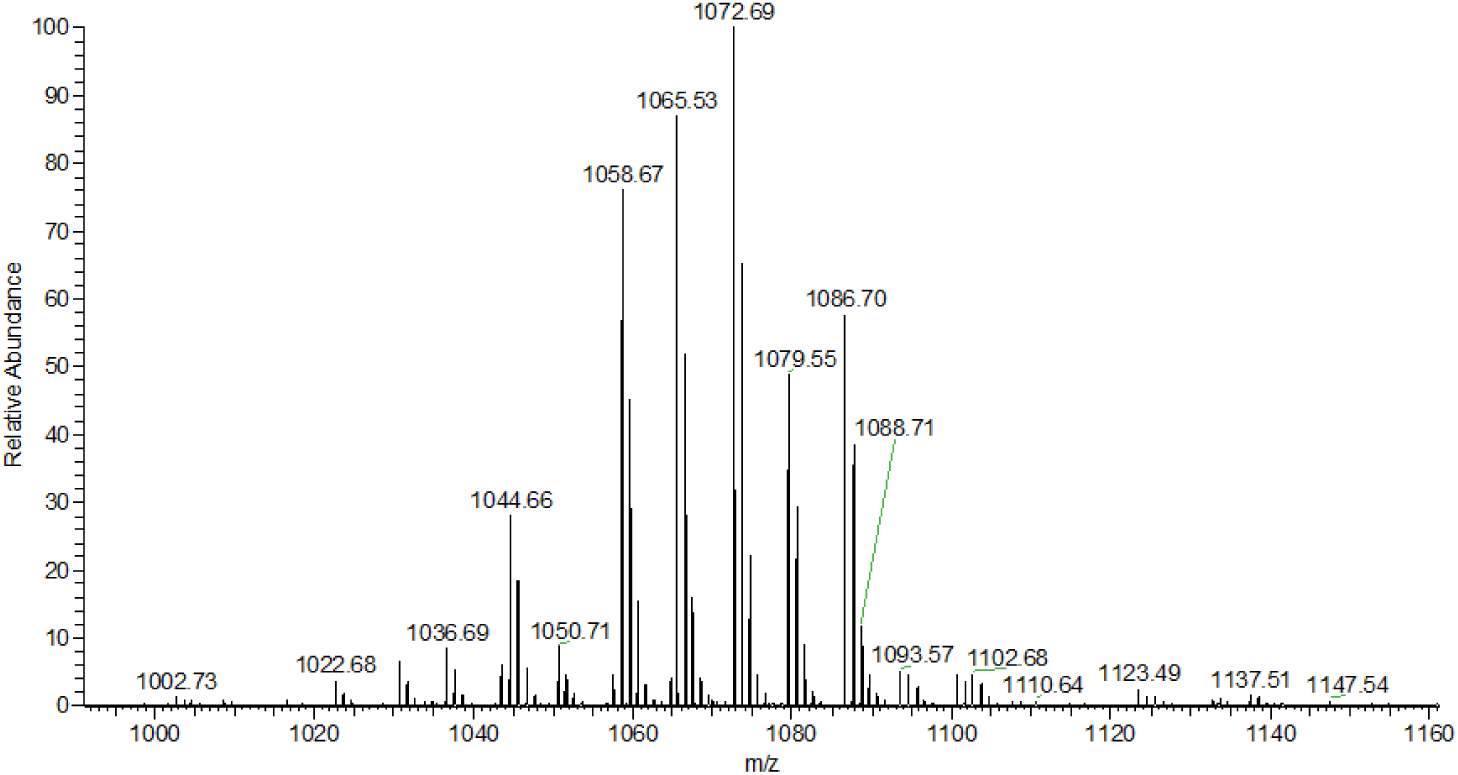
Analysis of CLE using MALDI-TOF-MS.

**Tab. 2.** Analysis of CLE using MALDI-TOF-MS.

| Lipopeptide | Fatty acid chain | Molecular formula | Molecular weight ($m/z$) | |
| --- | --- | --- | --- | --- |
|  |  |  | [M+H]<sup>+</sup> | [M+Na]<sup>+</sup> |
| Surfactin | C<sub>14</sub> | C<sub>52</sub>H<sub>91</sub>N<sub>7</sub>O<sub>13</sub> | 1,022.68 | 1,044.66 |
|  | C<sub>15</sub> | C<sub>53</sub>H<sub>93</sub>N<sub>7</sub>O<sub>13</sub> | 1,036.69 | 1,058.67 |
|  | C<sub>16</sub> | C<sub>54</sub>H<sub>95</sub>N<sub>7</sub>O<sub>13</sub> | 1,050.71 | 1,072.69 |
|  | C<sub>17</sub> | C<sub>55</sub>H<sub>97</sub>N<sub>7</sub>O<sub>13</sub> | - | 1,086.70 |
| IturinA | C<sub>14</sub> | C<sub>48</sub>H<sub>74</sub>N<sub>12</sub>O<sub>14</sub> | - | 1,065.53 |
|  | C<sub>15</sub> | C<sub>49</sub>H<sub>76</sub>N<sub>12</sub>O<sub>14</sub> | - | 1,079.55 |

### 2.5 Purification of CLE using HPLC system and FTIR analysis

Analysis of the retention time of commercial standard lipopeptides exhibited two obvious peaks at 21.6 min (peak c, surfactin) and 23.2 min (peak d, iturin). The corresponding peaks from the CLE group were collected, which were peak a at 21.4 min and peak b at 23.6 min (Fig. 7).

**Fig. 6.**
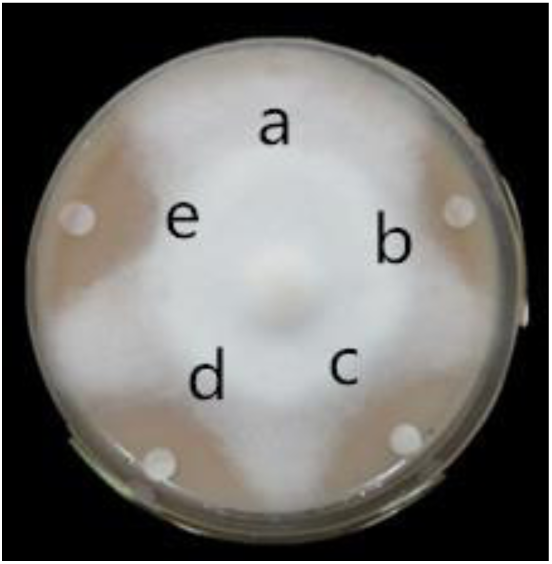
Inhibition effect of CLE on mycelial growth. Note: a: Control (distilled water); b: 1 mg/mL; c: 3 mg/mL; d: 5 mg/mL; e: Metalaxyl (15 µg/mL).

**Fig. 7.**
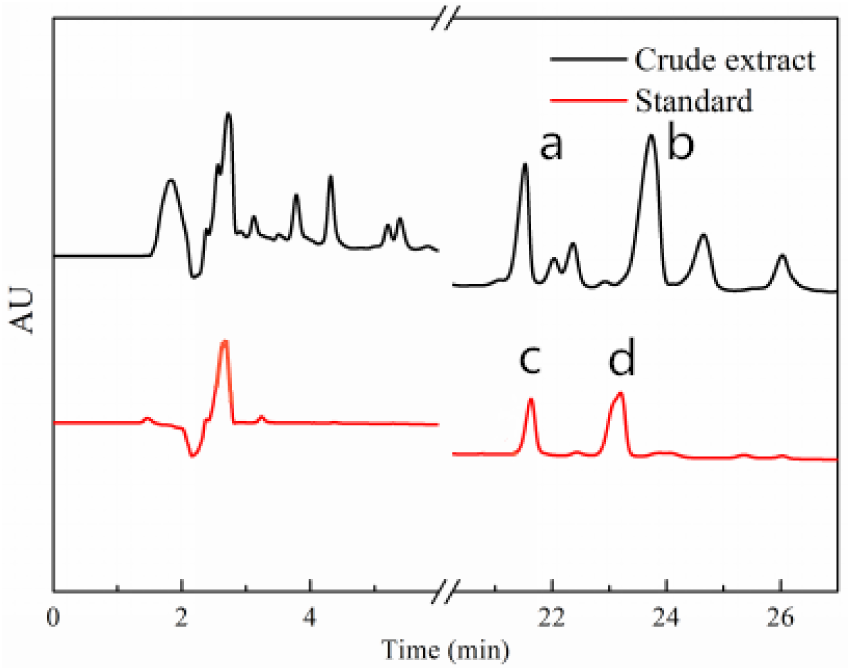
Purification of CLE using HPLC system. Note: The black line above shows the result for the CLE (peak a at 21.4 min, and peak b at 23.6 min). The red line below represents the results for standard lipopeptides; peak c at 21.6 min was commercial surfactin, and peak d at 23.2 min was commercial iturin.

From comparison of the results for purified surfactin and standard surfactin (Fig. 8A), a strong absorbance peak from 3,650 cm^-1^ to 3,250 cm^-1^ with a maximum at 3,292 cm^-1^ signified the presence of hydrogen-bonded -OH and -NH functional groups, which are characteristics of carbon-containing compounds with amino groups(74). Consecutive sharp absorbance peaks were found at 2,956, 2,925 and 2,854 cm^-1^, which correspond to the presence of -C-CH_3_ vibration banding or long alkyl chains(75). The highest peak at 1,664 cm^-1^ signified the presence of an amino acid zwitterion -C=O, which represented a peptide part(76). The weak absorbance peaks at 1,456 cm^-1^ and 1,406 cm^-1^ in the absorption signals ranging from 1,350 to 1,460 cm^-1^ were due to the -C-CH_2_ and -C-CH_3_ group vibrations contained in aliphatic chains(77). The peak at 1,194 cm^-1^ was probably due to the presence of C-O-C vibrations in esters(75, 78). The FTIR spectrum above showed that the combination of aliphatic groups with peptide moieties was a typical feature in lipopeptides. The comparison of results for purified IturinA and standard iturin (Fig. 8B) exhibited that the obvious peaks at 2,958, 2,925, 2,854, 1,458, and 1,386 cm^-1^ signified the aliphatic chains, and the peptide part was represented by the peaks at 3,307, 1,654, 1,541, and 1,205 cm^-1^.

**Fig. 8.**
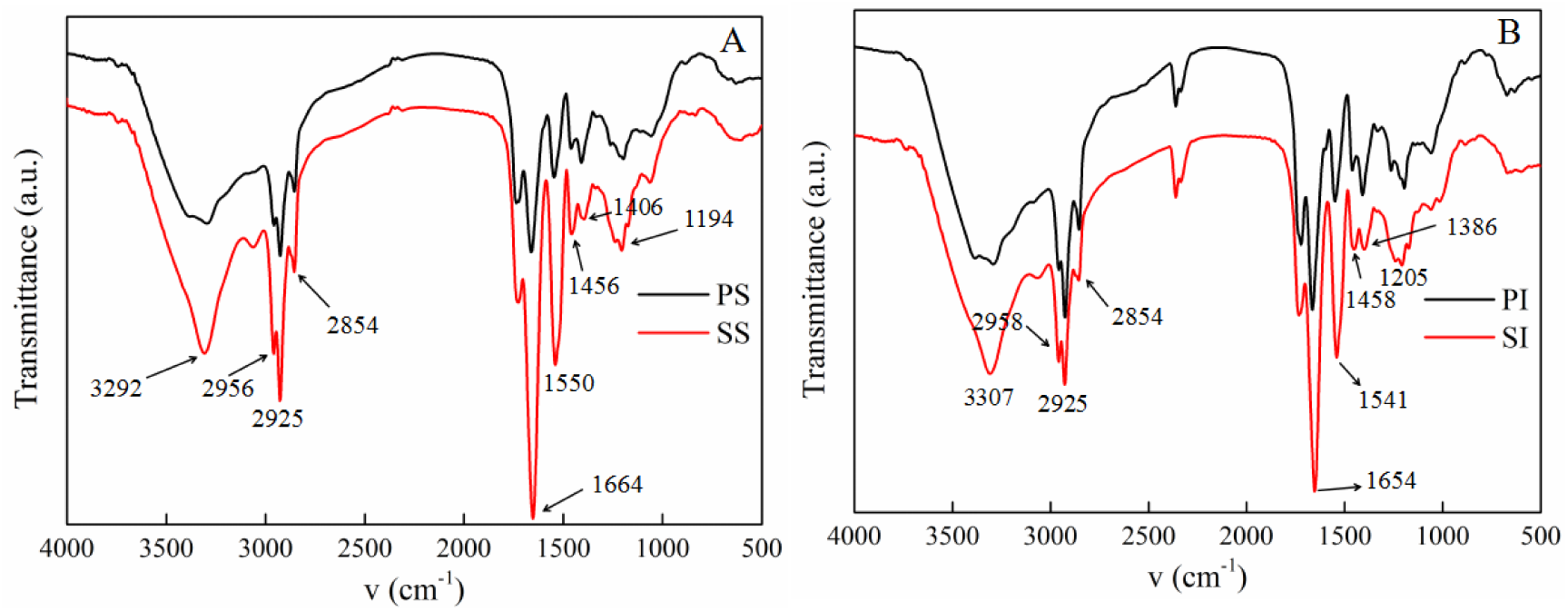
FTIR analysis of purified lipopeptides. Note: A: Comparison of purified surfactin (PS) and standard surfactin (SS); B: Comparison of purified IturinA (PI) and standard iturin (SI). The black line above shows the result for purified lipopeptides, and the red line below shows the result for standard lipopeptides.

### 2.6 MS/MS analysis of purified lipopeptides and comparison of their antioomycete activities

The fraction of peak a was subjected to MALDI-TOF-MS, and the *m/z* signals ranging from 1,000 to 1,100 were hypothesized to be produced by surfactin with fatty acid chains ranging in length from C_14_ to C_17_ (Fig. 9A, Tab. 4). In detail, the ion peaks at *m/z* 1,022.68, 1,044.66 and 1,060.68 were hypothesized to be the [M+H]^+^, [M+Na]^+^ and [M+K]^+^ adducts for surfactin C_14_ (1,022), and the ion peaks at *m/z* 1,036.69 [M+H]^+^, 1,058.67 [M+Na]^+^ and 1,074.64 [M+K]^+^ were assumed to be surfactin C_15_ (1,036). In addition, the ion peaks at *m/z* 1,050.71 [M+H]^+^, 1,072.69 [M+Na]^+^ and 1,088.66 [M+K]^+^ were considered surfactin C_16_ (1,050). Additionally, Na^+^ and K^+^ adduct ions of surfactin C_17_ (1,064) were deduced from *m/z* values of 1,086.69 and 1,102.68, respectively. Furthermore, MALDI-TOF-MS results of peak b with intense signals in the *m/z* range from 1,000 to 1,100 signified ions characteristic of IturinA C_14_ and IturinA C_15_ (Fig. 9B, Tab. 4). The peak series at *m/z* 1,043.55 [M+H]^+^, 1,065.53 [M+Na]^+^ and 1,081.56 [M+K]^+^ was suggestive of IturinA C_14_ (1,043), and Fig. 8B also showed ion peaks at *m/z* 1,057.57 [M+H]^+^, 1,079.55 [M+Na]^+^ and 1,093.56 [M+K]^+^, which all represented isoforms of IturinA C_15_ (1,057).

**Fig. 9.**
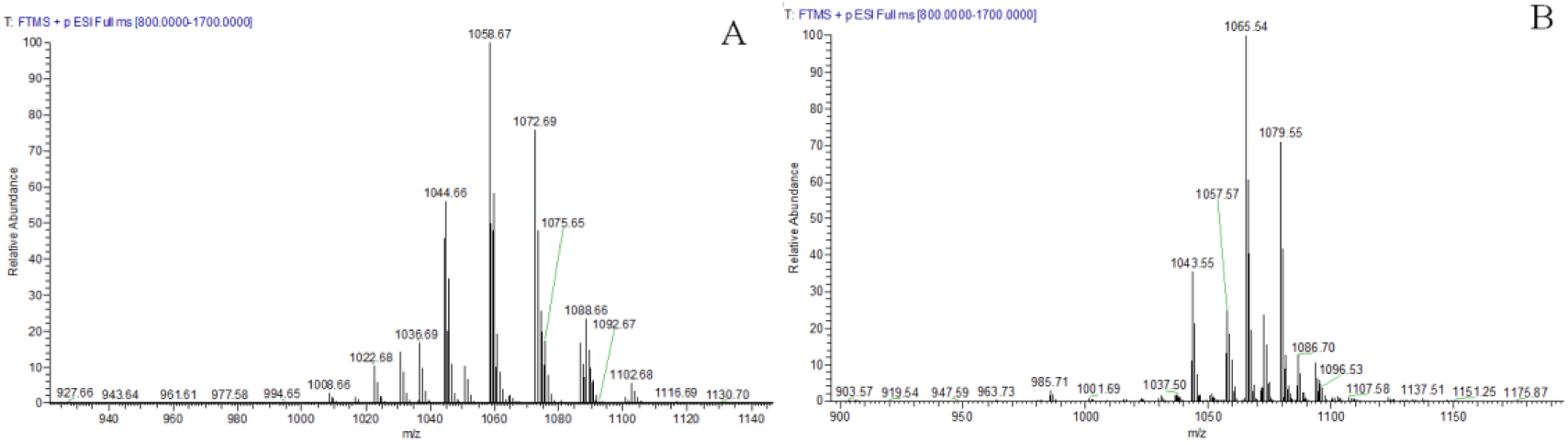
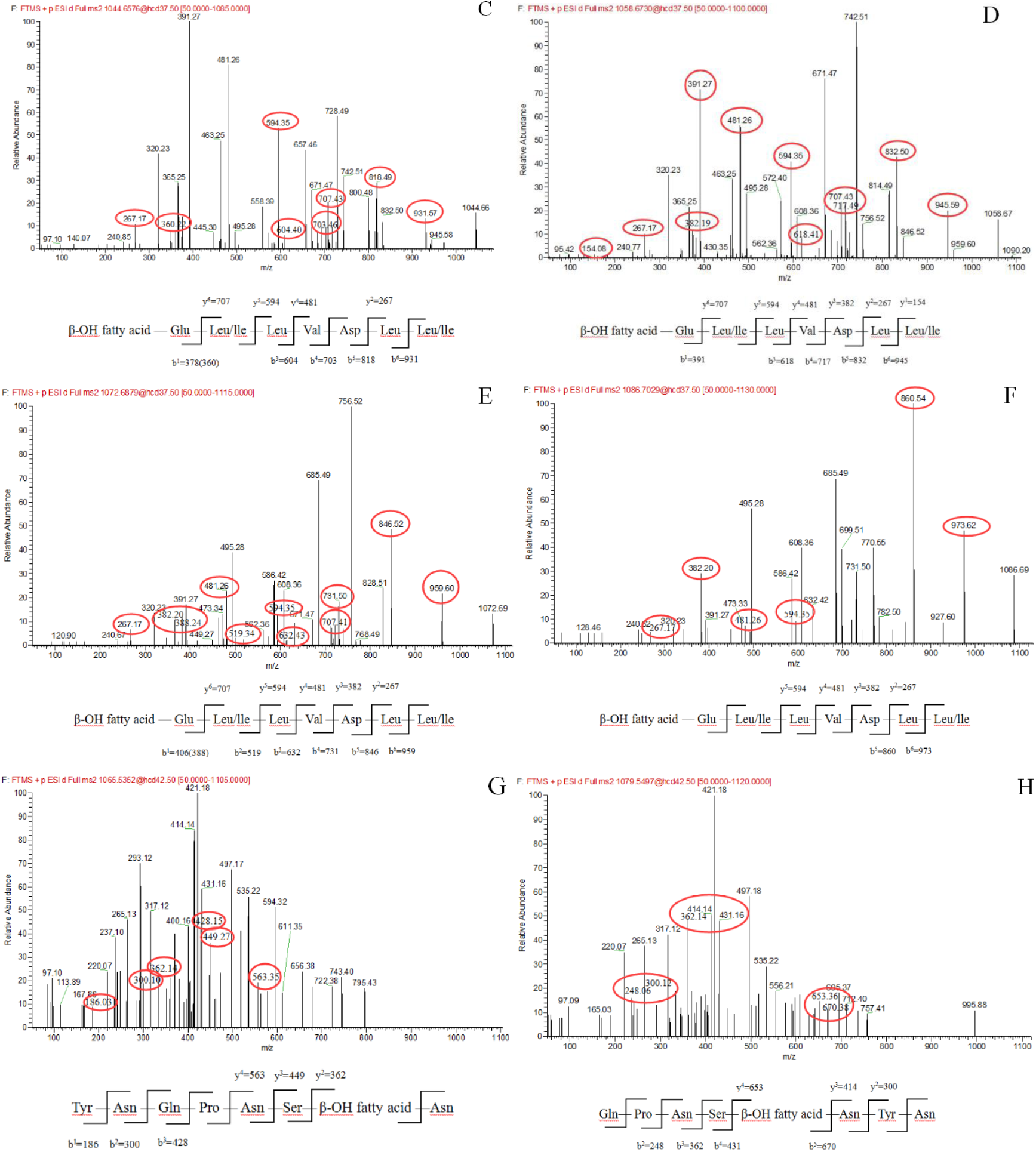
Detection of purified lipopeptides using MADI-TOF-MS/MS. Note: A: Full MS of peak a (surfactin); B: Full MS of peak b (IturinA); C-F: MS/MS spectrum of the surfactin C_14_ precursor ion at *m/z* 1,044.66 [M+Na]^+^, surfactin C_15_ precursor ion at *m/z* 1,058.67 [M+Na]^+^, surfactin C_16_ precursor ion at *m/z* 1,072.69 [M+Na]^+^ and surfactin C_17_ precursor ion at *m/z* 1,086.69 [M+Na]^+^, respectively; G-H: MS/MS spectrum of the IturinA C_14_ precursor ion at *m/z* 1,065.53 [M+Na]^+^ and spectrum of the IturinA C15 precursor ion at *m/z* 1,079.55 [M+Na]^+^.

**Tab. 3.**
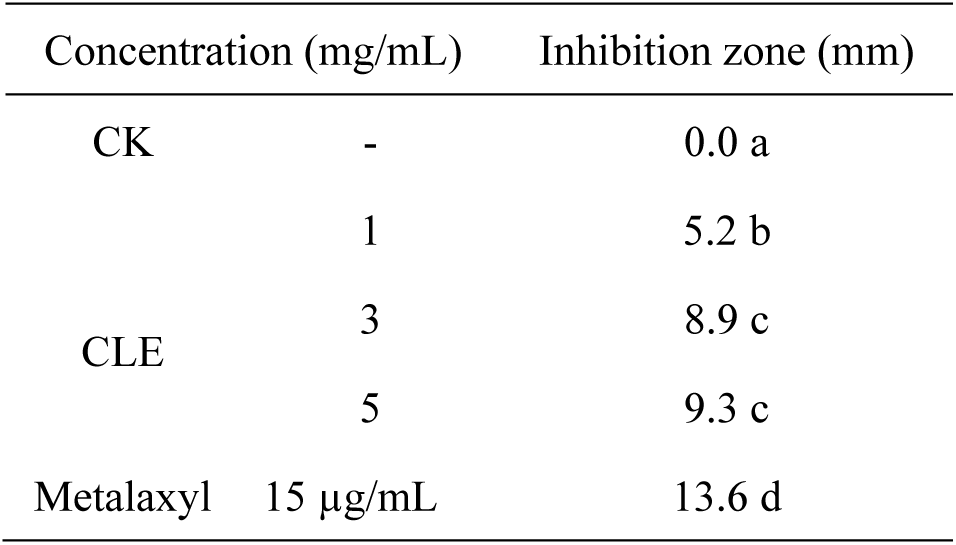
Inhibition effect of CLE on *P. infestans* mycelial growth.

| Concentration (mg/mL) |  | Inhibition zone (mm) |
| --- | --- | --- |
| CK | - | 0.0 a |
| CLE | 1 | 5.2 b |
|  | 3 | 8.9 c |
|  | 5 | 9.3 c |
| Metalaxyl | 15 µg/mL | 13.6 d |

**Tab. 4.** Detection of purified lipopeptides.

| Lipopeptide | Fatty acid chain | Molecular formula | Calculated (*m/z*) |  |  |
| --- | --- | --- | --- | --- | --- |
|  |  |  | [M+H]<sup>+</sup> | [M+Na]<sup>+</sup> | [M+K]<sup>+</sup> |
| surfactin<br>(peak a) | C<sub>14</sub> | C<sub>52</sub>H<sub>91</sub>N<sub>7</sub>O<sub>13</sub> | 1,022.68 | 1,044.66 | 1,060.68 |
|  | C<sub>15</sub> | C<sub>53</sub>H<sub>93</sub>N<sub>7</sub>O<sub>13</sub> | 1,036.69 | 1,058.67 | 1,074.64 |
|  | C<sub>16</sub> | C<sub>54</sub>H<sub>95</sub>N<sub>7</sub>O<sub>13</sub> | 1,050.71 | 1,072.69 | 1,088.66 |
|  | C<sub>17</sub> | C<sub>55</sub>H<sub>97</sub>N<sub>7</sub>O<sub>13</sub> | - | 1,086.69 | 1,102.68 |
| IturinA<br>(peak b) | C<sub>14</sub> | C<sub>48</sub>H<sub>74</sub>N<sub>12</sub>O<sub>14</sub> | 1,043.55 | 1,065.53 | 1,081.56 |
|  | C<sub>15</sub> | C<sub>49</sub>H<sub>76</sub>N<sub>12</sub>O<sub>14</sub> | 1,057.57 | 1,079.55 | 1,093.56 |

The amino acid sequences of the molecules of interest were detected using MS/MS. Fig. 9C illustrates the MS/MS spectrum of surfactin C_14_ at *m/z* 1,044.66 [M+Na]^+^. The series of b^+^ ions at *m/z* 931→818→703→604→378 (-H_2_O, 360) signified the loss of Leu, Asp, Val, and Leu-Leu/Ile at peptide bonds, and the ions at *m*/*z* 360 were the C terminus of a β-OH fatty acid combined with Glu. Starting from the y^+^ end, ions at *m/z* 267→481→594→707 represented the peptide bonds connected by Leu/Ile-Leu, Asp-Val, Leu, and Leu/Ile, respectively, so ions at *m/z* 707 were the total mass of ion fragments containing Leu/Ile-Leu-Val-Asp-Leu-Leu/Ile. The MS/MS spectrum exhibiting b^+^ and y^+^ fragment ions confirmed that the structure of surfactin C_14_ was β-OH fatty acid-Glu-Leu/Ile-Leu-Val-Asp-Leu-Leu/Ile. The structure of surfactin C_15_ at *m/z* 1,058.67 [M+Na]^+^ was determined by the result of Fig. 9D. Similar to above, the series of y^+^ ions at *m*/*z* 154→267→382→481→594→707 represented the connection of amino acids Leu/Ile, Leu, Asp, Val, Leu, and Leu/Ile, respectively. From the perspective of the b^+^ fragment, ions at *m/z* 945→832→717→618→391 illustrated the loss of Leu, Asp, Val, and Leu-Leu/Ile from the end of the C terminus, and the ions at *m*/*z* 391 were β-OH fatty acid connected with Glu. Meanwhile, the MS/MS spectrum of surfactin C_16_ at *m/z* 1,072.69 [M+Na]^+^ is represented in Fig. 9E. The set of y^+^ fragment ions was the same as those of surfactin C_14_ and surfactin C_15_, with the sequence of Leu/Ile-Leu-Val-Asp-Leu-Leu/Ile at the end of the N terminus. As a result of the b+ part, the most significant ion series at *m/z* 406 (-H_2_O, 388) confirmed the structure of β-OH fatty acid (C_16_) connected with Glu. Additionally, the y^+^ fragment ions that occurred in Na adducted ions, which were found at *m*/*z* 1,086.69 (C_17_, Fig. 9F), signified the same peptide connection in surfactin C_14-16_. The b^+^ fragment ions at *m/z* 973 explained the sequence of β-OH fatty acid (C_17_)-Glu-Leu/Ile-Leu-Val-Asp-Leu, and the ions at *m/z* 973→860 were the result of losing a Leu. In summary, the MS/MS spectrum peaks at *m*/*z* 1,044.66 (Fig. 9C), 1,058.67 (Fig. 9D), 1,072.69 (Fig. 9E), and 1,086.69 (Fig. 9F) were detected as the same subfamily (surfactin) but had a difference of 14 Da (-CH_2_-). The MS/MS spectrum of peak b (IturinA) is shown in Fig. 9G-H. IturinA at *m*/*z* 1,065.53 [M+Na]^+^ was analyzed in Fig. 9G. In detail, the b^+^ fragment ions at *m/z* 186→300→428 represented the sequence of Tyr-Asn-Gln, and ions at *m/z* 186 ions derived from a Tyr. In addition, the series of y^+^ ions at *m*/*z* 563→449→362 signified the cleavage and loss of Asn, Asp and Ser, respectively. The most significant ions at *m*/*z* 362 supported the fragment of β-OH fatty acid (C_14_)-Asn, and ions at *m/z* 563 illustrated the sequence of Asn-Ser-β-OH fatty acid-Asn. Fig. 9H shows the detection of IturinA at *m*/*z* 1,079.55 [M+Na]^+^. First, y^+^ fragment ions at *m*/*z* 300→414→653 symbolized the connection of Tyr-Asn, Asn and β-OH fatty acid, and ions at *m*/*z* 239 (653 - 414 = 239) matched exactly the fragment ion mass of β-OH fatty acid (C_15_). In addition, the b^+^ ion fragments in the order of *m*/*z* 248, 362, 431, and 670 illustrated the sequence of Gln-Pro-Asn-Ser-β-OH fatty acid. The results of Fig. 9G-H demonstrate IturinA C_14_ and IturinA C_15_ with a difference of 14 Da (-CH_2_-) and represent the structure of β-OH fatty acid-Asn-Tyr-Asn-Gln-Pro-Asn-Ser.

The antioomycete activity results (Fig. 10 and Tab. 5) showed that surfactin had no inhibition activity on the growth of mycelium and that there were no obvious inhibition zones at even the concentration of 50 µg/mL (Fig. 10A). However, the inhibitory effect was clearly dependent on the increasing concentration of IturinA (Fig. 10B). IturinA at the concentration of 50 µg/mL produced the best inhibition effect, and the inhibition zone reached a maximum of 10.5 mm (Fig. 10B-d). There was a significant difference between IturinA and the control (*P*<0.05).

**Fig. 10.**
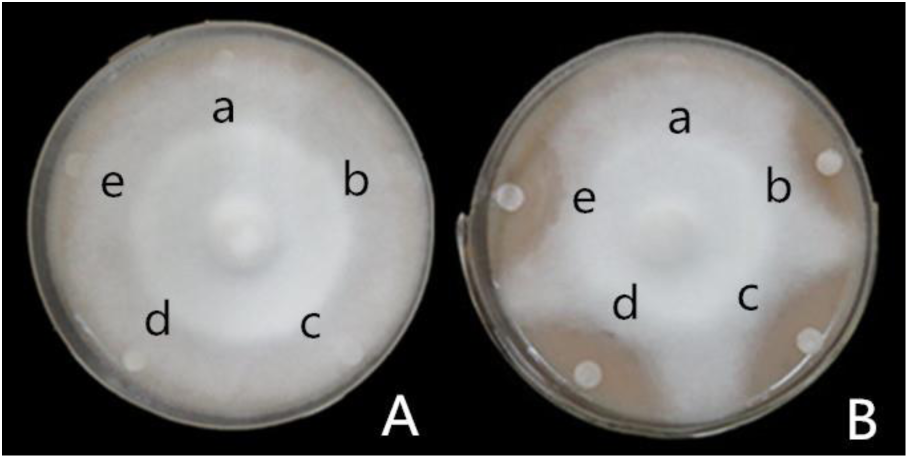
Inhibition effect of purified surfactin and IturinA. Note: A: Surfactin group; B: IturinA group. a: Control (distilled water); b: 20 µg/mL; c: 30 µ g/mL; d: 40 µg /mL; e: 50 µg/mL.

**Tab. 5.**
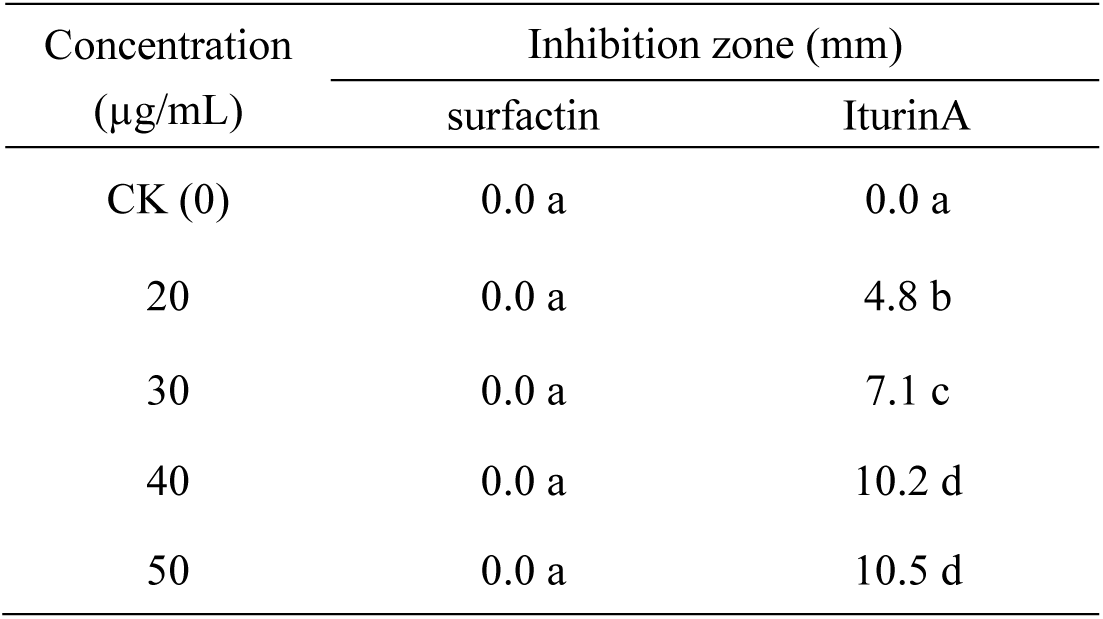
Inhibition effect of surfactin and IturinA against *P. infestans*.

### 2.7 Inhibition effect of IturinA against *P. infestans*

#### (1) The recovery of *P. infestans* mycelium and sporangium after inhibition

After inhibiting with IturinA (20, 30, 40, and 50 µg/mL), *P. infestans* mycelia recovered to grow (Fig. 11A, b-e) at a lower rate (7.1 mm/d, 5.3 mm/d, 3.2 mm/d, and 2.7 mm/d, respectively) than that of control (10.7 mm/d), and there were significant differences between IturinA treatments and the control (*P*<0.05). The results above signified that the concentration of IturinA was positively correlated with the degree of mycelium damage. After the inhibition of IturinA at the concentration of 50 µg/mL, the mycelium recovery rate exhibited the lowest value (only 2.7 mm/d, Fig. 11B). Meanwhile, zoospore release and sporangium direct germination rates were calculated, and the results showed that with the IturinA concentration increasing from 20 to 50 µg/mL, zoospore release and sporangium direct germination rates declined significantly. In detail, the zoospore release rate declined from 64.9% to 18.6%, and the sporangium direct germination rate decreased from 48.9% to 14.4% (Fig. 11C). The lowest zoospore release rate (18.6%) and sporangium direct germination rate (14.4%) that occurred after treatment with the highest IturinA concentration (50 µg/mL) were significantly different from those of the control (64.9% and 48.9%, respectively, *P*<0.05).

**Fig. 11.**
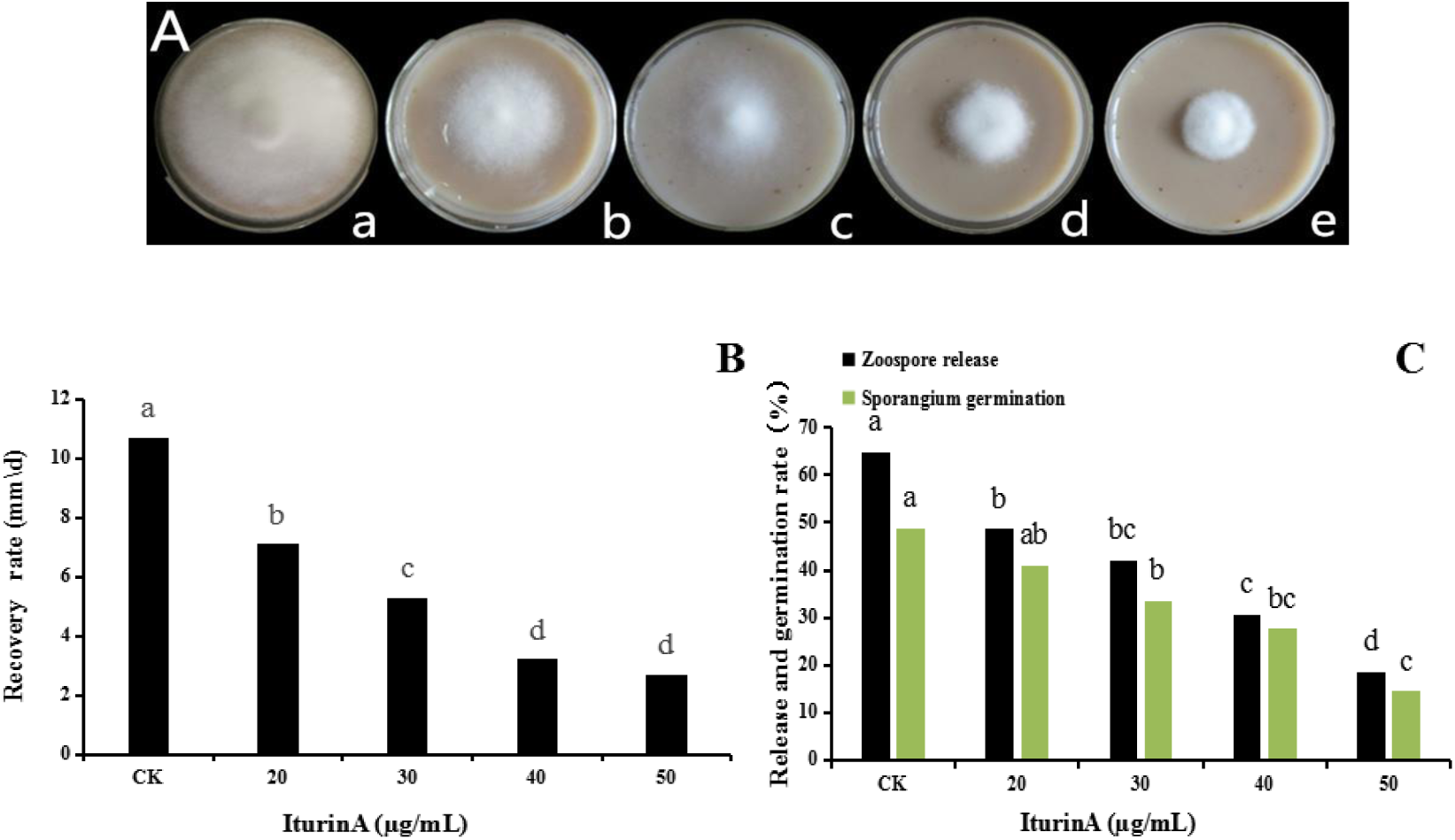
The recovery of *P. infestans* mycelia and sporangia after inhibition. Note: A-a: Control (normal mycelium growth); A-b: 20 µg/mL; A-c: 30 µg/mL; A-d: 40 µg/mL; A-e: 50 µg/mL. B: Mycelium recovery rate. C: Zoospore release and direct germination rates of sporangium after inhibition. The lowercase letters indicate a comparison between the different groups.

#### (2) Observation using OM, SEM and TEM

Under the OM examination, the mycelia in the control group (Fig. 12A-a) were smooth, vimineous, straight, and evenly grown. The mycelia affected by IturinA (50 µg/mL) exhibited a series of deformations (Fig. 12A, b-e). After treatment with IturinA, some mycelia twisted into clusters (Fig. 12A-b), some others grew with unequal widths, and abnormal branches were observed frequently (Fig. 12A-c). In addition, many mycelia lost smoothness and formed unusual surface bulges (Fig. 12A-d), the inner mycelium developed large vacuoles, and the cytoplasm condensed unevenly (Fig. 12A-e). An SEM system was used to observe mycelium deformation in shapes and appearances. The results showed that the mycelia in the control group were straight and smooth without any expansion (Fig. 12B-a). However, in the treatment group, mycelia were rough and uneven on the surface (Fig. 12B-b). In addition, mycelia were locally raised, with an uneven width (Fig. 12B-c,d). Expansion in branches (Fig. 12B-e) and even abnormal branches appeared in parts of the mycelium (Fig. 12B-f). The TEM method was used to examine the structural variation within cells. The results showed that normal mycelial cell membranes were intact, organelles within the cells were distributed in a normal arrangement, and mitochondria, including inner ridges, were abundant (Fig. 12C-a). After treatment with IturinA (50 µg/mL), the same mycelial cell membranes were disrupted (Fig. 12C, b-d), and organelles within the cell were disordered (Fig. 12C-b). A large area of cavitation appeared in the center of the cytoplasm (Fig. 12C-b,d), and mitochondria and ridges were sparse (Fig. 12C-b) compared with those of the control. Moreover, irregular organelle shapes with unclear boundaries and obvious accumulation bodies were also visible in some cells (Fig. 12C-c). Furthermore, organelles within some cells gathered in clumps, and the nuclei affected by cavitation shifted to the cell edge (Fig. 12C-d).

**Fig. 12.**
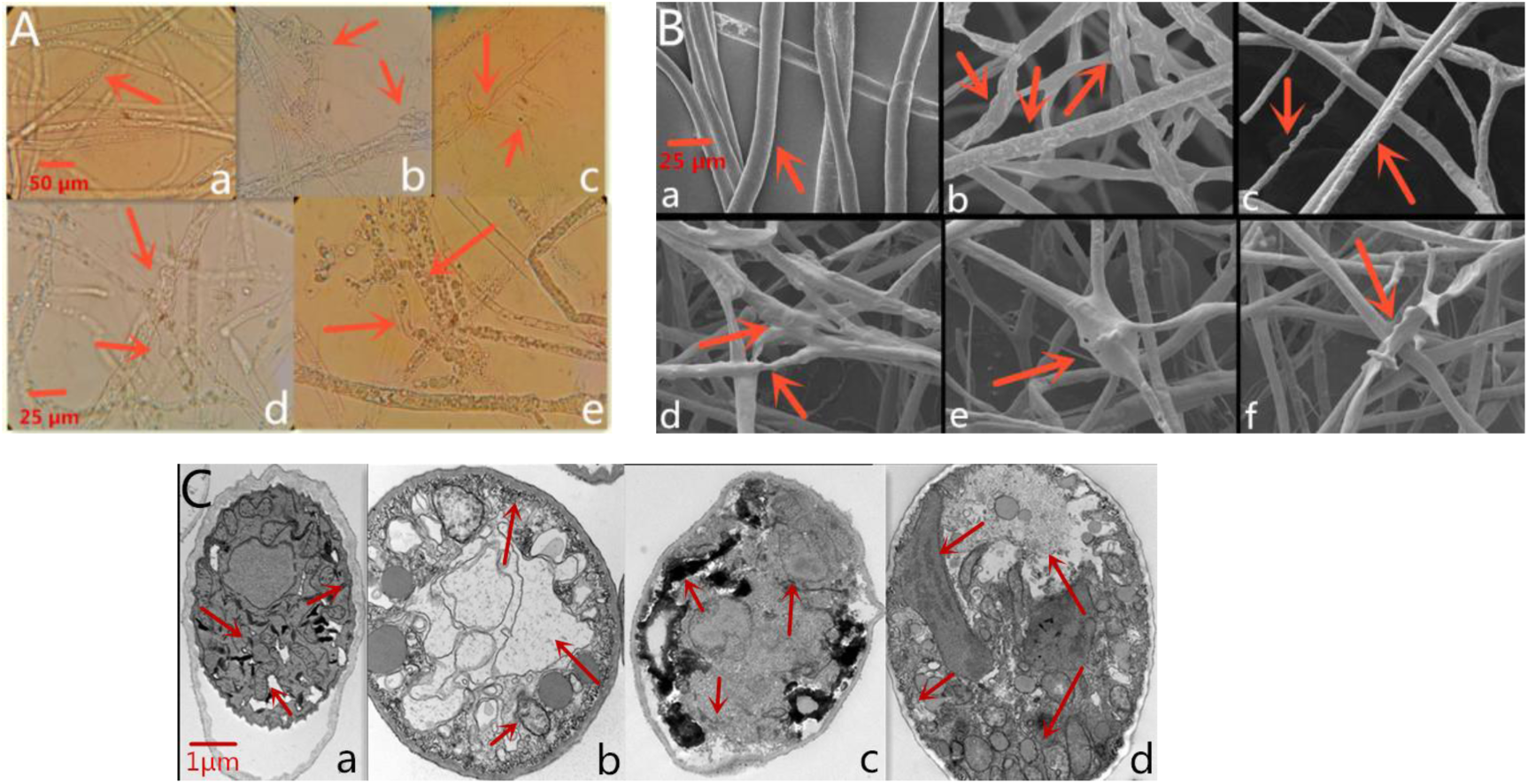
*P. infestans* mycelium deformation after inhibition. Note: A: OM observation (a-c, bar = 50 µm, and d-e, bar = 25 µm). A-a: Normal mycelium growth of the control, with a smooth, vimineous, straight and evenly grown mycelium; A-b: Mycelium twisted into clusters; A-c: Mycelium growth with unequal widths and increased branching; A-d: Loss of smoothness and formation of unusual surface bulges in the mycelium; A-e: Large vacuoles and condensed cytoplasm. B: SEM observation (bar = 25 µm). B-a: Straight and smooth mycelium (control group); B-b: Mycelium surface was rough and uneven; B-c,d: Mycelium was locally raised, with uneven width and roughness on the surface; B-e: Mycelium expansion of branches; B-f: Abnormal branches in the mycelium. C: TEM observation (bar = 1 µm). C-a: Mycelium grew normally, the mycelium cell membrane was intact, organelles were distributed in a normal arrangement, and there were numerous mitochondria with abundant inner ridges (control group); C-b: Disrupted cell membrane, disordered organelles, large cavitation area in the center, and sparse mitochondria with few ridges; C-c: Irregular organelles and body accumulation; C-d: Nonintact cell membrane, large cavitation area, organelles gathered in clumps and shifted nucleus.

#### (3) Effects of IturinA on the cell membrane

Cell membrane integrity of *P. infestans* mycelium were examined using propidium iodide. The results showed that after treatment with IturinA (50 µg/mL), hyphae (Fig. 13A-b) and sporangia (Fig. 13A-d) displayed obvious red fluorescence compared with those of the control (Fig. 13A-a, c), which indicated that IturinA could result in substantial cell membrane defects and cell death. Meanwhile, the red fluorescence rate that was exhibited in the sporangium was approximately 68% in the treatment group, while in the control group, the red fluorescence rate was lower and only 21%.

**Fig. 13.**
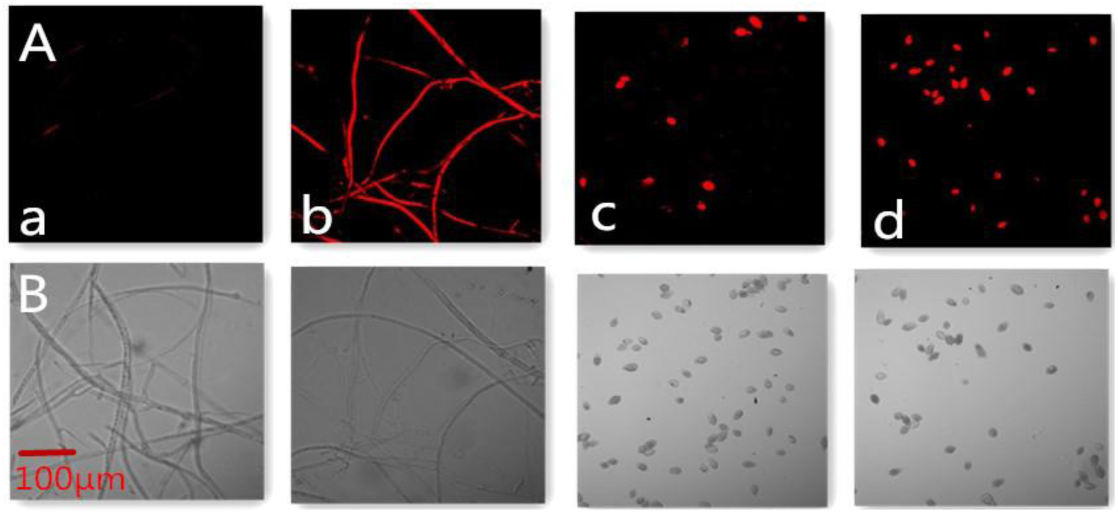
IturinA affects cell membrane integrity. Note: A: Observation in the red fluorescence channel; B: Observation in the optical channel. a-b, Mycelium; c-d, Sporangium. IturinA in the treatment group was at a concentration of 50 µg/mL.

The effect of IturinA on cell membrane permeability was shown in Fig. 14A. The relative conductivity of the control increased from 9.7% to 19.6% at 60 min. However, in the treatment (IturinA) group, relative conductivity improved from 10.2% at the beginning to 41.8%. In addition, the maximum relative conductivity of the treatment group (44.6%) was twice as high as that of the control group (20.9%). Leakage of nucleic acids revealed that at the soaking time of 100 min, the absorbance value reached a maximum of 0.251, which was significantly higher than that of the control group (the highest absorbance value was 0.059, Fig. 14B, *P*<0.05). In addition, detection of protein leakage showed that the highest absorbance value (0.410) appeared at 60 min, and the maximum absorbance of the treatment group was significantly higher than that of the control group (*P*<0.05), which had a maximum absorbance value of only 0.043 (Fig. 14C).

**Fig. 14.**
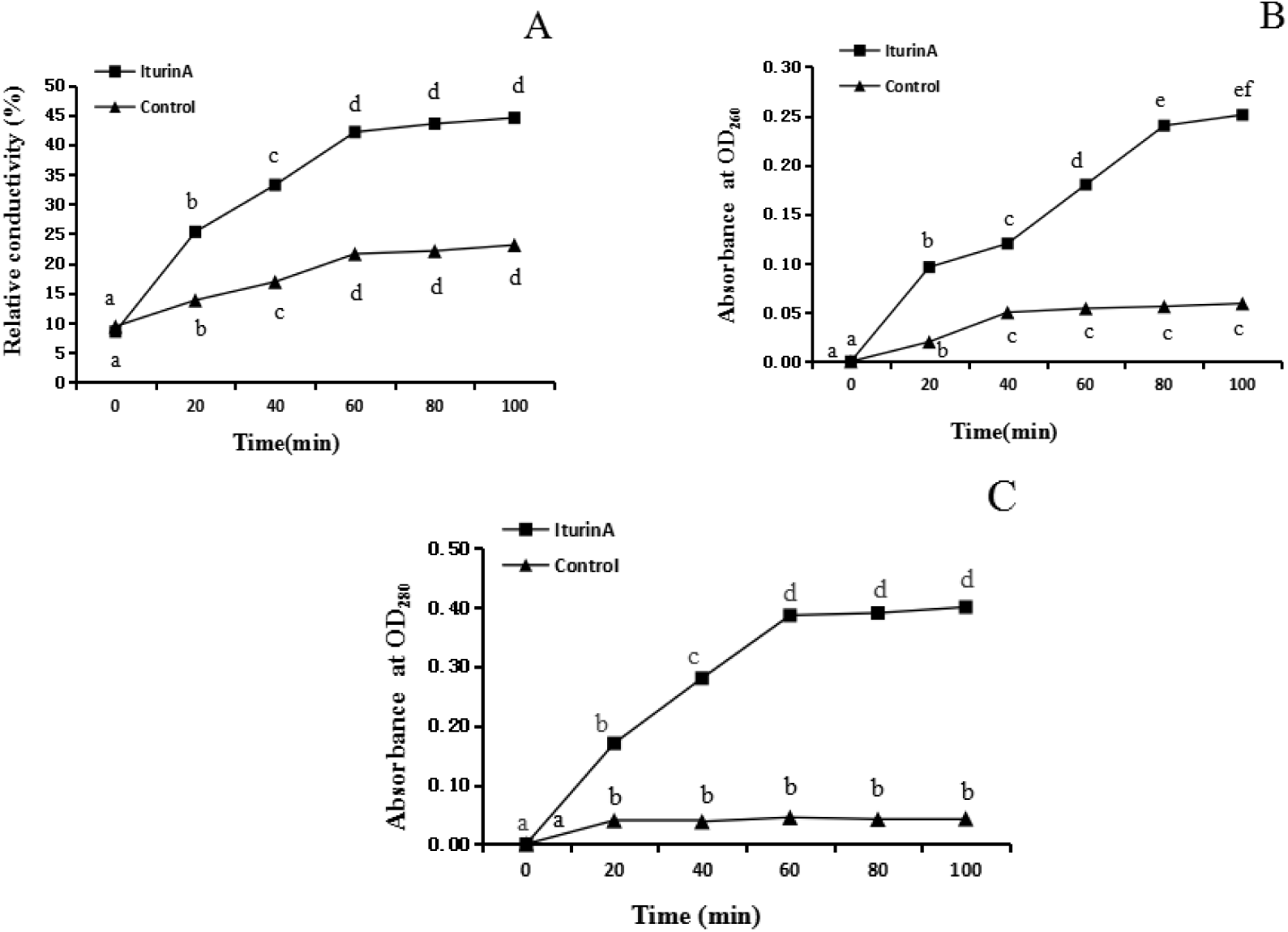
Effects of IturinA on cell membrane permeability. Note: A: Relative conductivity; B: Nucleic acid leakage; C: Protein leakage. IturinA in the treatment group was at a concentration of 50 µg/mL. The different letters indicate a significant difference (*P*<0.05).

### 2.8 ROS and MDA production

We hypothesized that IturinA application could lead to ROS generation, which is an important intermediate in the progression of *P. infestans* cell damage(79). While investigating this possibility, we observed a significant increase in intracellular ROS using DCFH-DA. As shown in Fig. 15, with increasing time of IturinA (50 µg/m L) treatment, the mean fluorescence intensity became obviously enhanced (Fig. 15A-B). In detail, the fluorescence intensity in treatment group was significantly higher than that in the control group after 4 h of generation (*P*<0.05). In the treatment group, the highest fluorescence intensity was four times higher than that in the control after 16 h of generation (*P*<0.05), and there was no significant difference between IturinA treatment and the positive control (Fig. 15A-B, *P*<0.05). In addition, the concentration of MDA produced by ROS reaction from 8 h to 24 h in the treatment group was significantly higher than that in the control group (approximately 20 µmol/L, *P*<0.05), and the highest MDA concentration reached a maximum of 152 µmol/L after 16 h of IturinA activity (Fig. 15C).

**Fig. 15.**
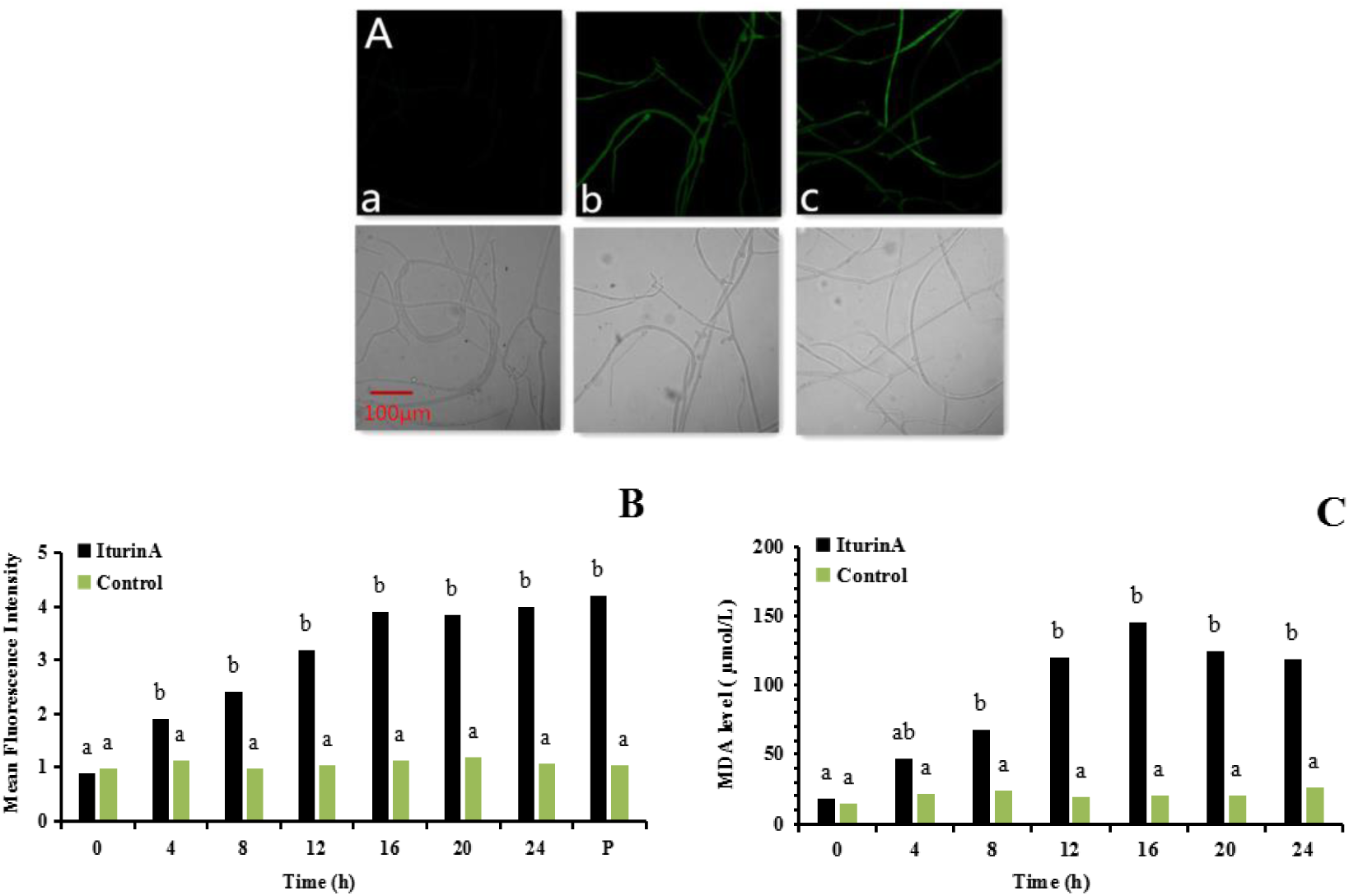
ROS generation and MDA production. Note: A: ROS detection. A-a: Control, A-b: IturinA (50 µg/mL), generation for 16 h, A-c: Positive control (P, Rosup), generation for 20 min. B: Mean fluorescence intensity. C: MDA production. The lowercase letters indicate a comparison within the same treatment group.

### 2.9 Mitochondrial damage

#### (1) Assay of MMP

The effect of IturinA on the MMP of *P. infestans* mycelium was detected using JC-1 staining and fluorescence microscopy. As shown in Fig. 16, the control group exhibited an obvious red fluorescence distribution (Fig. 16A-b) and J-aggregates (orange) in mitochondria (Fig. 16A-d). Compared with the control mycelia, IturinA-treated mycelia stained with JC-1 displayed dramatically changed fluorescence patterns and clear green fluorescence (Fig. 16B-c). These results indicated that IturinA could lead to a decrease in MMP.

**Fig. 16.**
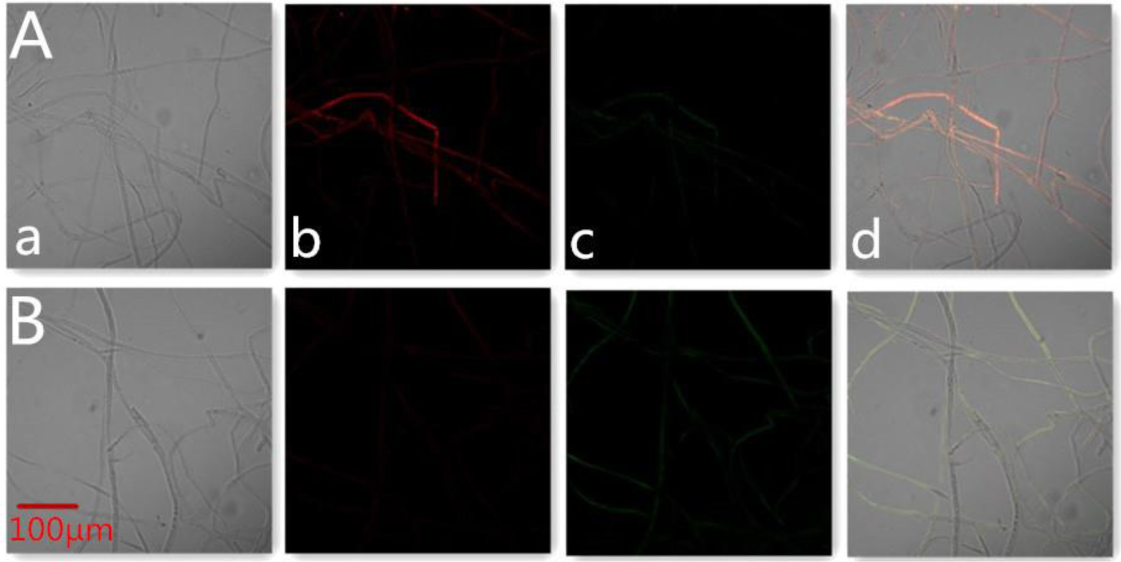
IturinA changes MMP. Note: A: Control group, B: Treatment group (IturinA). a: Optical channel; b: Red fluorescence channel; c: Green fluorescence channel; d: Red and green channels merged.

#### (2) MRCCA, RCR and P/O

The activities of complexes I-V were detected in this experiment, and the results are shown in Fig. 17A-E. Affected by IturinA, the activities of complex I-V respiratory enzymes were reduced remarkably and were approximately 61%, 35%, 43%, 31%, and 38%, respectively, which were significantly different from those of the control group (*P*<0.05). Meanwhile, the RCR and P/O values in the control group were 95% and 2.7, respectively, and in contrast, those in the treatment group were 63% and 1.9, respectively, which were significantly lower than those in the control group (Fig. 17F-G, *P*<0.05).

**Fig. 17.**
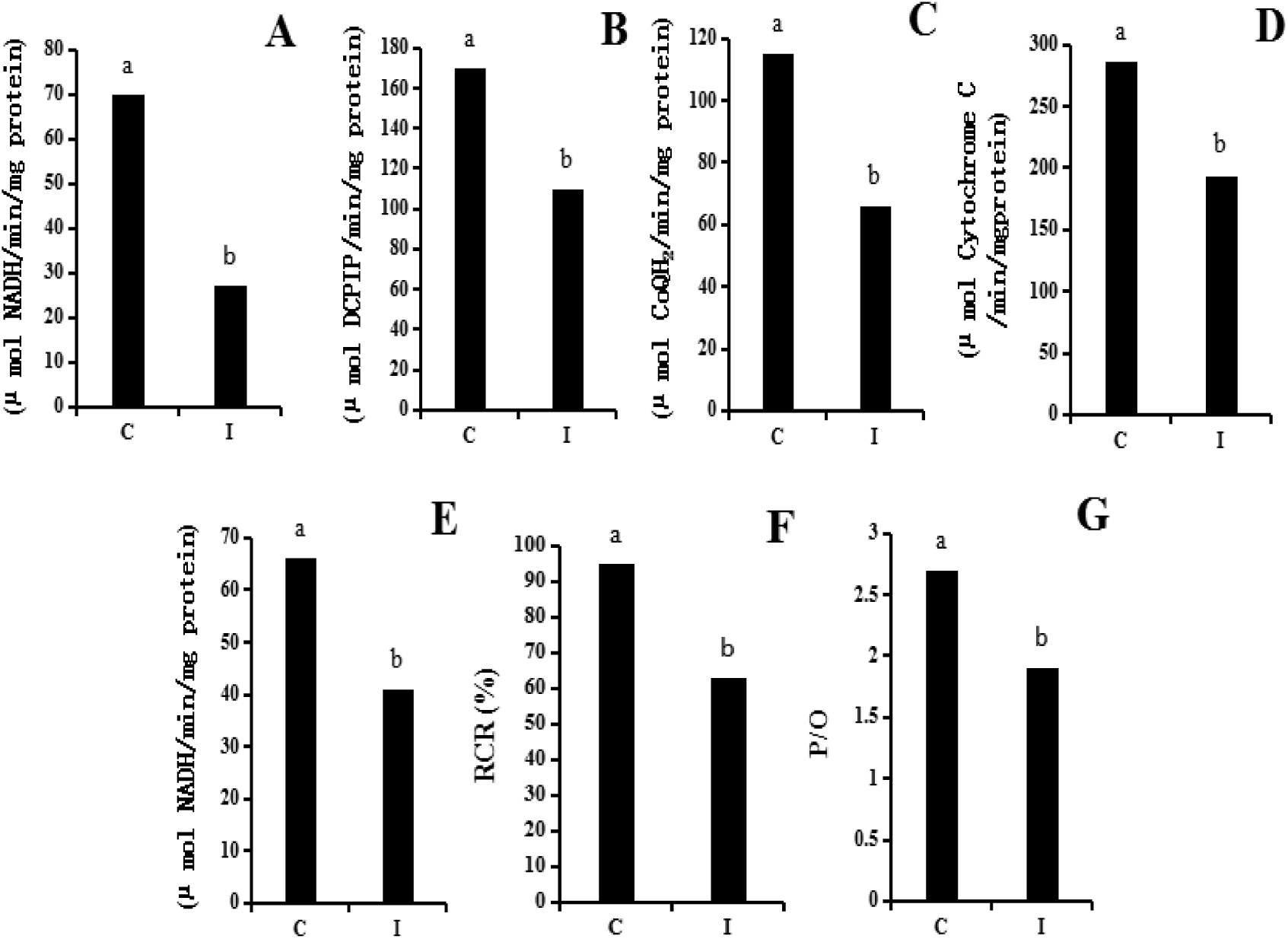
Detection of mitochondrial respiratory activity. Note: The letters C and I represent the control group and IturinA treatment group, respectively. A-E: Complex I to Complex V, respectively. F: Respiratory control rate (RCR). G: Oxidative phosphorylation efficiency (P/O).

## DISCUSSION

The increasing production of potatoes is still facing significant losses because of the infection of fungi, oomycetes, bacteria, insects, and viruses(80, 81). Among these pathogens, the *P. infestans* oomycete is the culprit of potato late blight, which is the disease that is the most serious and has the largest economic loss(1). The control of potato late blight based on the massive use of BCAs, including microorganisms and secondary metabolites, could be a potential measure to relieve or overcome the problem of food safety, environmental protection and disease resistance resulting from chemicals(7). Some *Bacillus* and *Pseudomonas* species are considered the best potential candidates used as BCAs because of their diversity, survival ability in various environments, and their variety of biocontrol molecules(82, 83). Additionally, as the result of massive number of bioactive compounds involved in their antagonistic activity, *Bacillus* and *Pseudomonas* species also have numerous interesting properties for industry and agriculture(84). In this study, we compared the inhibitory effects of LC, CS, and CFS from three bacterial species, *B. subtilis* WL-2, *P. fluorescens* WL-1, and *B. pumilus* W-7, against *P. infestans* mycelium growth. The inhibition effect of WL-2 was overall significantly better than that of the other strains. Although the biocontrol strains exhibited a strong antifungal effect in a plate confrontation test, the biological control effect in the *in vitro* experiment was still worrying due to the environmental changes(85). Based on previous experience, we suspect that antagonistic strains themselves are destructive to potato tissues(86). Therefore, in this study, the WL-2 strain was selected to test the effect of controlling potato late blight on tissues *in vitro*. The results indicated that the WL-2 strain had an obvious ability to prevent late blight development on tissues *in vitro* and had no side effect on potato tissues.

In fact, CLPs with a wide range of antibacterial activities are some of the most abundant and highly yielded metabolites from *Bacillus*(9). In addition, the peptide cycle with 7 ∼ 10 amino acids combined with a lipid component (β-hydroxy fatty acid chain or β-amino fatty acid chain) determines that CLPs are amphiphilic compounds(3). Furthermore, the hemolysis, oil dispersal, and emulsification activities(34, 37, 38) could be preliminarily detected to demonstrate the capability of CLPs secretion. The obvious transparent trace (Fig. 3B-a), large oil dispersal ring (diameter = 4.92 cm), high percentage of emulsification (82.1%), and decline in ST indicated that the WL-2 strain had a strong ability to produce CLPs. With the character that CLPs aggregate and precipitate at the condition of pH = 2, the acid precipitation method(41) was used to prepare CLE. CFS obtained from *Bacillus* species possesses various bioactive substances, such as polysaccharides, proteins, lipids, and peptides(87); however, whether CLE has the ability to inhibit *P. infestans* mycelium growth should be further investigated. In this study, our results showed that when the CLE concentration was 5 mg/mL, the obvious inhibition zone reached a maximum of 9.3 mm.

Based on the obvious inhibitory effect of various homologous subfamilies contained in CLPs, it was extremely meaningful to determine the CLPs classification and clarify the molecular mechanism of *P. infestans* inhibition. In this part, by comparison with standard lipopeptides, we showed the same retention time via HPLC detection and the same absorption peaks pattern in FTIR analysis, demonstrating that both subfamilies of surfactin and IturinA were presented in the CLE. Additionally, in the FTIR spectrum, the aliphatic groups observed at 2,958, 2,925, 2,854, 1,458, and 1,386 cm^-1^ were connected with the peptide parts exhibited at 3,307, 1,654, 1,541, and 1,205 cm^-1^, indicating that the purified CLPs of surfactin and IturinA possess an amphiphilic trait(75, 77, 78). Further study was conducted using MS/MS technology to detect the specific molecular weight and structural formula according to the amino acid numbers and sequence(16). The results showed that the chemical structural formula of purified surfactin was β-OH fatty acid-Glu-Leu/Ile-Leu-Val-Asp-Leu-Leu/Ile with a fatty acid chain from C_14_ to C_17_. The purified IturinA with ions characteristic of IturinA C_14_ and IturinA C_15_ had a structure of β-OH fatty acid-Asn-Tyr-Asn-Gln-Pro-Asn-Ser. However, based on previous results, *B. amyloliquefaciens* S76-3 could produce the CLPs PlipastatinA and IturinA(51); *B. amyloliquefaciens* PGPBacCA1 could produce surfactin, IturinA and fengycin(88), and *B. subtilis* BS155 has the ability to produce surfactin and fengycin(17). We found that the CLP types produced by different antagonistic strains exhibited a great diversity. In addition, environmental factors, such as pathogens, temperature, and carbon and nitrogen sources, could also affect the classification, production and proportion of different CLP subfamilies to change the antagonistic effect(49, 89).

Iturin and fengycin produced by *Bacillus* spp. are known to exhibit direct antifungal activity(83, 90, 91), and their fungal toxicity mechanisms are involved in pore formation in the cell membrane(9, 92). Similarly, a CLP with 9 amino acids produced by *P. fluorescens* SBW25 has direct antioomycete activities and results in immobilization and subsequent lysis of *P. infestans* zoospores(93, 94). However, the specific inhibitory effects of surfactin and iturin against *P. infestans* remain unclear. After defining the types of CLPs in this research, the inhibition mechanisms of IturinA and surfactin against *P. infestans* were the most important issues to explore in this research.

Subsequently, our results showed that surfactin had no direct inhibition activity on the growth of *P. infestans* mycelium (Fig. 10A), which is similar to reports that surfactin alone lacks antifungal activities(14, 17, 89). However, some research results indicated that the surfactin family produced by *Bacillus* spp. has an indirect antagonistic activity by triggering induced systemic resistance (ISR) in plant(95–97). This indirect activity of surfactin on potato plants against late blight should be investigated as a meaningful work in the future. Most interesting to us was the fact that after direct inhibition with IturinA (50 µg/mL), the inhibition zone against *P. infestans* mycelium growth reached a maximum of 10.5 mm (Fig. 10B-e), and the lowest zoospore release and sporangium direct germination rate were only 18.6% and 14.4%, respectively. These results corresponded to the report that the fengycin family produced by *P. fluorescens* SBW25 has a specific inhibitory effect on *P. infestans* zoospore activity through zoospore membrane solubilization(94). In fact, the inhibited *P. infestans* mycelium must be in a damaged state; however, the specific injury mechanisms caused by IturinA remain unclear. Just as former articles reported that Iturin produced by *Bacillus* species exhibited direct fungal toxicity involving cell membrane damage and pore formation in the plasma membrane(9, 92), in this research, we found that the affected *P. infestans* mycelia were rough and uneven on the surface (Fig. 12B-b), and that unusual surface bulges (Fig. 12A-d) were formed in the mycelia. Much of the changes in mycelial appearance are probably due to the damage of the internal cell structure(53, 54). Next, our TEM results showed that the inhibited cell membranes were disrupted (Fig. 12C, b-d), organelles adopted an irregular shape (Fig. 12C-c) and were disordered (Fig. 12C-b), and a large area of cavitation appeared in the center of the cytoplasm (Fig. 12C-b,d). All the OM, SEM and TEM analysis results were basically similar to previous findings, with a report that *F. graminearum* mycelium affected by IturinA derived from *B. amyloliquefaciens* S76-3 displayed severe morphological changes, including mycelium distortions, cell membrane leakage, and separation of the plasma membrane from the cell wall(51). In contrast with reported cellular content inactivation and branch formation inhibition(51), in our research, many tiny and irregular branches (Fig. 12A-c) stretched to the surrounding environment to evade the toxic effects of IturinA.

Moreover, the cell membrane damage probably led to the release of nucleic acids and protein from the cell and directly changed the relative conductivity of the mycelium-soaked solution(60). Additionally, the fluorescent dye propidium iodide is a kind of nucleus-staining reagent, and the red fluorescence displayed by propidium iodide can distinguish damage of the cell membrane from an intact membrane present in a living state (17). In this part, our results showed that IturinA results in *P. infestans* mycelium cell membrane defects and cell death, that the inhibited hyphae (Fig. 13A-b) and sporangia (Fig. 13A-d) displayed red fluorescence, and that the ratio of sporangia with red fluorescence reached a maximum of 68%. In addition, when inhibited by IturinA, the released protein and nucleic acids increased the relative conductivity of the mycelium-soaked solution to approximately two times higher than that of the control (Fig. 14).

Intracellular chaos caused by long-term adversity could also induce ROS generation in cells to adapt to the adverse environment(54). The ROS generation caused by detrimental conditions is an important intermediate in the progression of cell damage(62). In our research, ROS detection results showed that after 4 h of exposure to IturinA, the highest ROS generation was four times as high as that of the control. In addition, the highest MDA concentration reached a maximum of 152 µmol/L after 16 h of treatment. However, possibly because MDA is a subsequent product of ROS generation, the highest values of ROS generation and MDA concentration did not appear at the same time(60, 98). In addition, when living in harsh environments and affected by ROS generation, mitochondria might develop an abnormal state, in which the cell respiratory process is obstructed, and the power plants needed for cell life might have abnormal working conditions(63, 70–72). During ROS generation, the accumulation of oxidized products could also lead to MRCCA decline and electron transport chain dysfunction resulting in an immature respiration process, which ultimately leads to a decrease in P/O(65, 70–72).

In this study, JC-1 staining showed that IturinA leads to a decrease in MMP in *P. infestans* cells. The respiratory enzyme activity of complexes I-V declined by approximately 61%, 35%, 43%, 31%, and 38%, respectively (Fig. 17A-E). Meanwhile, the RCR and P/O values were only 63% and 1.9, respectively, which were significantly lower than those of the control (Fig. 17F-G, *P*<0.05). Energy production is closely related to mitochondrial function and oxidative phosphorylation processes(80). Therefore, the decline in MMP and MRCCA in *P. infestans* and the weakness of ATP production in *P. infestans* mitochondria strongly indicated that IturinA resulted in serious mitochondrial damage that affected cellular respiratory state. Taken together, these data clarified that the WL-2 strain can produce the CLPs surfactin and IturinA. Surfactin had no direct inhibitory effect on *P. infestans* mycelium growth, While IturinA could cause *P. infestans* cell membrane disruption, induce cellular ROS generation and, most importantly, lead to mitochondrial damage, blocking ATP production. All the results above highlight that *B. subtilis* WL-2 and its IturinA lipopeptides have great potential for inhibiting *P. infestans* mycelium growth and controlling the development of potato late blight in the future.

In this article, we have performed many studies on controlling potato late blight using CLPs; however, many issues are worth resolving. For instance, the indirect inhibition effects and the differences among surfactin, Iturin, and fengycin in triggering the ISR in potato plants are still unknown. The inducer from pathogens aimed at CLPs seems to be specific for one of the CLPs subfamilies, for example, *Fusarium oxysporum* significantly induced fengycin production by *B. amyloliquefaciens* SQR9, while when strain SQR9 was induced by other pathogens (*Rhizoctonia solani* and *Fusarium solani*), surfactin production increased obviously, and fengycin secretion decreased significantly(99). Therefore, this specific inducing phenomenon with a potentially high impact for biological control is well worth knowing in future. Cooperation of surfactin with iturin or fengycin is still a controversial issue. Parent Zihalirwa Kulimushi once suggested that surfactin from *B. amyloliquefaciens* FZB42 could somehow interfere with fengycin activity against *Rhizomucor variabilis*(89). Additionally, a mixture the CLPs surfactin and fengycin against *Verticillium dahlia* and *Rhizopus stolonifer* also lost the inhibitory effect of fengycin on spore germination and hyphal growth(100, 101). This phenomenon may be explained by the stabilizing effect of surfactin on certain lipid bilayers(101, 102) and by the inactive complexes formed by coaggregation of surfactin and fengycin(103). In contrast, the cooperation of surfactin with iturin and fengycin extracted from *Bacillus velezensis* (Y6 and F7) against *Ralstonia solanacearum* and *F. oxysporum* displayed an obviously improved antifungal effect(61). Therefore, the relationship of surfactin and iturin regarding inhibition of *P. infestans* should also be investigated in future research.

## ACKNOWLEDGMENTS

This research was supported by the Agriculture Special Scientific Research Program of China (grant No. 201303018 to Jizhi Jiang), the Natural Science Foundation Program of China (grant No. C11474083 to Yanqing Wu), the Natural Science Foundation Program of Hebei Province of China (grant No. C2015201231 to Jizhi Jiang), and the Hebei University Postgraduate Innovation Program (grant No. X2016073 to Youyou Wang).

## REFERENCES

1. Schepers HTAM, Kessel GJT, Lucca F, Förch MG, van den Bosch GBM, Topper CG, Evenhuis A. 2018. Reduced efficacy of fluazinam against *Phytophthora infestans* in the Netherlands. Eur J Plant Pathol 151:947–960.

2. Dey T, Saville A, Myers K, Tewari S, Cooke DEL, Tripathy S, Fry WE, Ristaino JB, Guha Roy S. 2018. Large sub-clonal variation in *Phytophthora infestans* from recent severe late blight epidemics in India. Sci Rep 8:4429.

3. Wang YY. 2018. The study of antagonistic bacteria WL2 against Phytophthora infestans and its lipopeptides on disease prevention and growth promotion. Hebei University.

4. Jin GH, Li XZ, Wang YC, Wang T. 2017. Effects of inter-annual drought on the complexity of physiological races of *Phytophthora infestans*. Plant Protection 43:167–173.

5. Fukue Y, Akino S, Osawa H, Kondo N. 2018. Races of *Phytophthora infestans* isolated from potato in Hokkaido, Japan. J Gen Plant Pathol 84:276–278.

6. Bajwa R, Khalid A, Cheema TS. 2003. Antifungal activity of allelopathic plant extracts III: growth response of some pathogenic fungi to aqueous extract of *Parthenium hysterophorus*. Plant Pathol J 2:145–156.

7. Raj MK, Kanwar SS. 2015. Lipopeptides as the antifungal and antibacterial agents: applications in food safety and therapeutics. Biomed Res Int 2015:473050.

8. Kim HJ, Choi HS, Yang SY, Kim IS, Yamaguchi T, Sohng JK, Park SK, Kim JC, Lee CH, Gardener BM, Kim YC. 2014. Both extracellular chitinase and a new cyclic lipopeptide, chromobactomycin, contribute to the biocontrol activity of *Chromobacterium* sp. C61. Mol Plant Pathol 15:122–132.

9. Ongena M, Jacques P. 2008. *Bacillus* lipopeptides: versatile weapons for plant disease biocontrol. Trends Microbiol 16:115–125.

10. Aranda FJ, Teruel JA, Ortiz A. 2005. Further aspects on the hemolytic activity of the antibiotic lipopeptide iturin A. Biochim Biophys Acta 1713:51–56.

11. Zhang QX, Zhang Y, Shan HH, Tong YH, Chen XJ, Liu FQ. 2017. Isolation and identification of antifungal peptides from *Bacillus amyloliquefaciens* W10. Environ Sci Pollut Res Int 24:25000–25009.

12. Tsuge K, Akiyama T, Shoda M. 2001. Cloning, sequencing, and characterization of the IturinA operon. J Bacteriol 183:6265–6273.

13. Alvarez F, Castro M, Príncipe A, Borioli G, Fischer S, Mori G, Jofré E. 2015. The plant-associated *Bacillus amyloliquefaciens* strains MEP218 and ARP23 capable of producing the cyclic lipopeptides iturin or surfactin and fengycin are effective in biocontrol of sclerotinia stem rot disease. J Appl Microbiol 112:159–174.

14. Ben Abdallah D, Frikha-Gargouri O, Tounsi S. 2015. *Bacillus amyloliquefaciens* strain 32a as a source of lipopeptides for biocontrol of *Agrobacterium tumefaciens* strains. J Appl Microbiol 119:196–207.

15. Zeriouh H, de Vicente A, Perez-Garcia A, Romero D. 2014. Surfactin triggers biofilm formation of *Bacillus subtilis* in melon phylloplane and contributes to the biocontrol activity. Environ Microbiol 16:2196–211.

16. Yang H, Li X, Li X, Yu H, Shen Z. 2015. Identification of lipopeptide isoforms by MALDI-TOF-MS/MS based on the simultaneous purification of iturin, fengycin, and surfactin by RP-HPLC. Anal Bioanal Chem 407:2529–2542.

17. Zhang L, Sun C. 2018. Cyclic lipopeptides fengycins from marine bacterium *Bacillus subtilis* kill plant pathogenic fungus *Magnaporthe grisea* by inducing reactive oxygen species production and chromatin condensation. Appl Environ Microbiol 84 :e00445–18.

18. Moyne AL, Shelby R, Cleveland TE, Tuzun S. 2001. Bacillomycin D: an iturin with antifungal activity against *Aspergillus flavus*. J Appl Microbiol 90:622–629.

19. Kumar A, Saini S, Wray V, Nimtz M, Prakash A, Johri BN. 2012. Characterization of an antifungal compound produced by *Bacillus* sp. strain A5F that inhibits *Sclerotinia sclerotiorum*. J Basic Microbiol 52:670–678.

20. Tabbene O, Di Grazia A, Azaiez S, Ben Slimene I, Elkahoui S, Alfeddy MN, Casciaro B, Luca V, Limam F, Mangoni ML. 2015. Synergistic fungicidal activity of the lipopeptide bacillomycin D with amphotericin B against pathogenic *Candida* species. FEMS Yeast Res 15:fov022.

21. Kefi A, Ben Slimene I, Karkouch I, Rihouey C, Azaeiz S, Bejaoui M, Belaid R, Cosette P, Jouenne T, Limam F. 2015. Characterization of endophytic *Bacillus* strains from tomato plants (*Lycopersicon esculentum*) displaying antifungal activity against *Botrytis cinerea* Pers. World J Microbiol Biotechnol 31:1967–1976.

22. Gu Q, Yang Y, Yuan Q, Shi G, Wu L, Lou Z, Huo R, Wu H, Borriss R, Gao X. 2017. Bacillomycin D produced by *Bacillus amyloliquefaciens* is involved in the antagonistic interaction with the plant-pathogenic fungus *Fusarium graminearum*. Appl Environ Microbiol 83:e01075–17.

23. Arrebola E, Jacobs R, Korsten L. 2010. Iturin A is the principal inhibitor in the biocontrol activity of *Bacillus amyloliquefaciens* PPCB004 against postharvest fungal pathogens. J Appl Microbiol 108:386–95.

24. Wang YY, Jiang JZ, Li Y, Zhang YH, Sun H, Lang YF. 2017. Inhibition comparison of six antagonistic bacteria against *Phytophthora infestans*. Journal of Hebei University(Natural Science Edition) 37:169–175.

25. Ali GS, El-Sayed AS, Patel JS, Green KB, Ali M, Brennan M, Norman D. 2015. *Ex Vivo* application of secreted metabolites produced by soil-inhabiting *Bacillus* spp. efficiently controls foliar diseases caused by *Alternaria* spp. Appl Environ Microbiol 82:478–490.

26. Bayston K, Tomlinson M, Cohen J. 1992. In-vitro stimulation of TNF-α from human whole blood by cell-free supernatants of Gram-positive bacteria. Cytokine 4:397–402.

27. Ndlovu T, Rautenbach M, Vosloo JA, Khan S, Khan W. 2017. Characterisation and antimicrobial activity of biosurfactant extracts produced by *Bacillus amyloliquefaciens* and *Pseudomonas aeruginosa* isolated from a wastewater treatment plant. AMB Express 7:108.

28. Huang YJ, Jiang JZ, Feng LN, Tian Y, Zhao S. 2014. Inhibition comparison of several antagonists against *Phytophthora infestans*. Journal of Hebei Agricultural University 37:80–85.

29. Kunova A, Bonaldi M, Saracchi M, Pizzatti C, Chen XY, Cortesi P. 2016. Selection of *Streptomyces* against soil borne fungal pathogens by a standardized dual culture assay and evaluation of their effects on seed germination and plant growth. BMC Microbiol 16:272.

30. Ding T, Su B, Chen X, Xie S, Gu S, Wang Q, Huang D, Jiang H. 2017. An endophytic bacterial strain isolated from *Eucommia ulmoides* inhibits southern corn leaf blight. Front Microbiol 8:903.

31. Jiang JZ, Liang TY, Wang HY, Wang XZ. 2013. Screening of antagonistic *Pseudomonas Fluorescens* against *Phytophthora infestans* and disease control *in vitro*. Journal of Agricultural University of Hebei 36:72–76.

32. Jiang JZ, Wang YY, Wang XN, Li LY, Wan AQ, Li M. 2017. Identification of SR13-2 strain against *Phytophthora infestans* and control of late blight on detached potato tissues. Crops 2017:146–150.

33. Balint-Kurti PJ, Zwonitzer JC, Wisser RJ, Carson ML, Oropeza-Rosas MA, Holland JB, Szalma SJ. 2007. Precise mapping of quantitative trait loci for resistance to southern leaf blight, caused by *Cochliobolus heterostrophus* race O, and flowering time using advanced intercross maize lines. Genetics 176:645–657.

34. Wu YQ, Wang YY, Wang C, Cha MY. 2018. Inhibitory effect of lipopeptide crude extract produced by *Bacillus subtilis* WL2 on *Phytophthora infestans* and its isolation and identificatication. Journal of Hebei University(Natural Science Edition) 38:632–639.

35. Biswas SC, Dubreil L, Marion D. 2001. Interfacial behavior of wheat puroindolines: study of adsorption at the air-water interface from surface tension measurement using wilhelmy plate method. J Coll Interf Sci 244:245–253.

36. Ceresa C, Rinaldi M, Chiono V, Carmagnola I, Allegrone G, Fracchia L. 2016. Lipopeptides from *Bacillus subtilis* AC7 inhibit adhesion and biofilm formation of *Candida albicans* on silicone. Antonie Van Leeuwenhoek 109:1375–1388.

37. Morikawa M, Hirata Y, Imanaka T. 2000. A study on the structure-function relationship of lipopeptide biosurfactants. Biochim Biophys Acta 14:211–218.

38. Huang W. 2011. Studies on degradation of oil wastewater by biosurfactant- producing bacteria. Journal of Hunan Agricultural University(Natural Sciences) 37:461–464.

39. Sen S, Borah SN, Bora A, Deka S. 2017. Production, characterization, and antifungal activity of a biosurfactant produced by *Rhodotorula babjevae* YS3. Microb Cell Fact 16:95.

40. Landy M, Warren GH, RosenmanM SB, Colio LG. 1948. Bacillomycin: an antibiotic from *Bacillus subtilis* active against pathogenic fungi. Exp Biol Med 67:539–541.

41. Chen X, Zhang Y, Fu X, Li Y, Wang Q. 2016. Isolation and characterization of *Bacillus amyloliquefaciens* PG12 for the biological control of apple ring rot. Postharvest Biol Technol 115:113–121.

42. Jiang J, Gao L, Bie X, Lu Z, Liu H, Zhang C, Lu F, Zhao H. 2016. Identification of novel surfactin derivatives from NRPS modification of *Bacillus subtilis* and its antifungal activity against *Fusarium moniliforme*. BMC Microbiol 16:31.

43. Bauer AW, Kirby WM, Sherris JC, Turck M. 1966. Antibiotic susceptibility testing by a standardized single disk method. Am J Clin Pathol 45:493–496.

44. Perez KJ, Viana JD, Lopes FC, Pereira JQ, Dos Santos DM, Oliveira JS, Velho RV, Crispim SM, Nicoli JR, Brandelli A, Nardi RM. 2017. *Bacillus* spp. isolated from puba as a source of biosurfactants and antimicrobial lipopeptides. Front Microbiol 8:61.

45. Fan H, Zhang Z, Li Y, Zhang X, Duan Y, Wang Q. 2017. Biocontrol of bacterial fruit blotch by *Bacillus subtilis* 9407 via surfactin-mediated antibacterial activity and colonization. Front Microbiol 8:1973.

46. Simionato AS, Navarro MOP, de Jesus MLA, Barazetti AR, da Silva CS, Simoes GC, Balbi-Pena MI, de Mello JCP, Panagio LA, de Almeida RSC, Andrade G, de Oliveira AG. 2017. The effect of phenazine-1-carboxylic acid on mycelial growth of *Botrytis cinerea* produced by *Pseudomonas aeruginosa* LV strain. Front Microbiol 8:1102.

47. Jha SS, Joshi SJ, S JG. 2016. Lipopeptide production by *Bacillus subtilis* R1 and its possible applications. Braz J Microbiol 47:955–964.

48. Jemil N, Ben Ayed H, Manresa A, Nasri M, Hmidet N. 2017. Antioxidant properties, antimicrobial and anti-adhesive activities of DCS1 lipopeptides from *Bacillus methylotrophicus* DCS1. BMC Microbiol 17:144.

49. Parthipan P, Preetham E, Machuca LL, Rahman PK, Murugan K, Rajasekar A. 2017. Biosurfactant and degradative enzymes mediated crude oil degradation by bacterium *Bacillus subtilis* A1. Front Microbiol 8:193.

50. Asari S, Ongena M, Debois D, De Pauw E, Chen K, Bejai S, Meijer J. 2017. Insights into the molecular basis of biocontrol of *Brassica* pathogens by *Bacillus amyloliquefaciens* UCMB5113 lipopeptides. Ann Bot 120:551–562.

51. Gong AD, Li HP, Yuan QS, Song XS, Yao W, He WJ, Zhang JB, Liao YC. 2015. Antagonistic mechanism of iturin A and plipastatin A from *Bacillus amyloliquefaciens* S76-3 from wheat spikes against *Fusarium graminearum*. PLoS One 10:e0116871.

52. Wang YY, Jiang JZ, Li M, Wang XN, Wu YQ. 2017. Comparative study on inhibition of several antagonistic bacteria against spore germination of *Phytophthora infestans*. China Plant Protection 37:16–23.

53. Cui ZN, Li YS, Hu DK, Tian H, Jiang JZ, Wang Y, Yan XJ. 2016. Synthesis and fungicidal activity of novel 2,5-disubstituted-1,3,4-thiadiazole derivatives containing 5-phenyl-2-furan. Sci Rep 6:20204.

54. Huiskonen JT. 2018. Image processing for cryogenic transmission electron microscopy of symmetry-mismatched complexes. Biosci Rep 38:BSR 20170203.

55. Tang H, Chen W, Dou ZM, Chen R, Hu Y, Chen W, Chen H. 2017. Antimicrobial effect of black pepper petroleum ether extract for the morphology of *Listeria monocytogenes* and *Salmonella typhimurium*. J Food Sci Technol 54:2067–2076.

56. Zhao P, Quan C, Wang Y, Wang J, Fan S. 2014. *Bacillus amyloliquefaciens* Q-426 as a potential biocontrol agent against *Fusarium oxysporum* f. sp. *spinaciae*. J Basic Microbiol 54:448–56.

57. Li X, Zhang Y, Wei Z, Guan Z, Cai Y, Liao X. 2016. Antifungal activity of isolated *Bacillus amyloliquefaciens* SYBC H47 for the biocontrol of peach gummosis. PLoS One 11:e0162125.

58. Cui Y, Zhao Y, Tian Y, Zhang W, Lü X, Jiang X. 2012. The molecular mechanism of action of bactericidal gold nanoparticles on *Escherichia coli*. Biomaterials 33:2327–2333.

59. Tian F, Li B, Ji B, Zhang G, Luo Y. 2009. Identification and structure-activity relationship of gallotannins separated from *Galla chinensis*. LWT Food Sci Technol 42:1289–1295.

60. Dolezalova E, Lukes P. 2015. Membrane damage and active but nonculturable state in liquid cultures of *Escherichia coli* treated with an atmospheric pressure plasma jet. Bioelectrochemistry 103:7–14.

61. Kobayashi D, Kondo K, Uehara N, Otokozawa S, Tsuji N, Yagihashi A, Watanabe N. 2002. Endogenous reactive oxygen species is an important mediator of miconazole antifungal effect. Antimicrob Agents Ch 46:3113–3117.

62. Tian J, Gan Y, Pan C, Zhang M, Wang XY, Tang XD, Peng X. 2018. Nerol-induced apoptosis associated with the generation of ROS and Ca^2+^ overload in saprotrophic fungus *Aspergillus flavus*. Appl Microbiol Biotechnol 102:6659.

63. Salvioli S, Ardizzoni A, Franceschi C, Cossarizza A. 1997. JC-1, but not DiOC_6_(3) or rhodamine 123, is a reliable fluorescent probe to assess delta psi changes in intact cells: implications for studies on mitochondrial functionality during apoptosis. FEBS Lett 411:77–82.

64. Pushpanathan M, Gunasekaran P, Rajendhran J. 2013. Mechanisms of the antifungal action of marine metagenome-derived peptide, MMGP1, against *Candida albicans*. PLoS One 8:e69316.

65. Sun MT, Mei W, Zi JJ, Yan CB, Chen Y, Yu M, Xiong W. 2019. Effect of MTERF2 on mitochondrial oxidative phosphorylation activity in human cervical cancer HeLa cells. Med & Pharm J Chin PLA 31:50–55.

66. Srere PA. 1969. Citrate synthase. Methods Enzymol 13:3–11.

67. Hermann Schägger, Pfeiffer K. 2001. The ratio of oxidative phosphorylation complexes I-V in bovine heart mitochondria and the composition of respiratory chain supercomplexes. J Biol Chem 276:37861–37867.

68. Davies SMK, Poljak A, Duncan MW, Smythe G A, Murphy MP. 2001. Measurement of protein carbonyls, *ortho-*and *meta-*tyrosine and oxidative phosphorylation complex activity in mitochondria from young and old rats. Free Radic Biol Med 31:181–190.

69. James AM, Wei YH, Pang CY, Murphy MP. 1996. Altered mitochondrial function in fibroblasts containing MELAS or MERRF mitochondrial DNA mutations. Biochem J 318:401–407.

70. Amaroli A, Ravera S, Baldini F, Benedicenti S, Panfoli I, Vergani L. 2019. Photobiomodulation with 808-nm diode laser light promotes wound healing of human endothelial cells through increased reactive oxygen species production stimulating mitochondrial oxidative phosphorylation. Lasers Med Sci 34:495–504.

71. Villani G, Attardi G. 2007. Polarographic assays of respiratory chain complex activity*. Methods Cell Biol 80:121–133.

72. Hinkle PC. 2004. P/O ratios of mitochondrial oxidative phosphorylation. Biochim Biophys Acta 1706:1–11.

73. Daverey A, Pakshirajan K. 2009. Production, characterization, and properties of sophorolipids from the yeast *Candida bombicola* using a low-cost fermentative medium. Appl Biochem Biotechnol 158:663–674.

74. Kong J, Yu S. 2007. Fourier transform infrared spectroscopic analysis of protein secondary structures. Acta Biochim Biophys Sin (Shanghai) 39:549–559.

75. Thaniyavarn J, Roongsawang N, Kameyama T, Haruki M, Imanaka T, Morikawa M, Kanaya S. 2003. Production and characterization of biosurfactants from *Bacillus licheniformis* F2.2. Biosci Biotechnol Biochem 67:1239–1244.

76. Kiran GS, Priyadharsini S, Sajayan A, Priyadharsini GB, Poulose N, Selvin J. 2017. Production of lipopeptide biosurfactant by a *Marine Nesterenkonia* sp. and its application in food industry. Front Microbiol 8:1138.

77. Dehghan-Noudeh G, Housaindokht M, Bazzaz BS. 2005. Isolation, characterization, and investigation of surface and hemolytic activities of a lipopeptide biosurfactant produced by *Bacillus subtilis* ATTC 6633. J Microbiol 43:272–276.

78. Das P, Mukherjee S, Sen R. 2008. Antimicrobial potential of a lipopeptide biosurfactant derived from a marine *Bacillus circulans*. J Appl Microbiol 104:1675–1684.

79. Madeo F, Frohlich E, Ligr M, Grey M, Sigrist SJ, Wolf DH, Frohlich KU. 1999. Oxygen stress: a regulator of apoptosis in yeast. J Cell Biol 145:757–767.

80. Lin Lf, Liu YL, FU S, Qu CH, Li H, Ni J. 2019. Inhibition of mitochondrial complex function-the hepatotoxicity mechanism of emodin based on quantitative proteomic analyses. Cells 8:263.

81. Alkher H, Islam MR, Wijekoon C, Kalischuk M, Kawchuk LM, Peters RD, Al-Mughrabi KI, Conn KL, Dobinson KF, Waterer D, Daayf F. 2015. Characterization of *phytophthora infestans* populations in canada during 2012. Can J Plant Pathol 37:305–314.

82. Kloepper JW, Reddy MS, Kenney DS, Vavrina C, Kokalis-Burelle N, Martinez-Ochoa N. 2004. Application for rhizobacteria in transplant production and yield enhancement. Int Soc Hortic Sci 631:219–229.

83. Raaijmakers JM, De Bruijn I, Nybroe O, Ongena M. 2010. Natural functions of lipopeptides from *Bacillus* and *Pseudomonas*: more than surfactants and antibiotics. FEMS Microbiol Rev 34:1037–1062.

84. Caulier S, Gillis A, Colau G, Licciardi F, Liépin M, Desoignies N, Modrie P, Legrève A, Mahillon J, Bragard C. 2018. Versatile antagonistic activities of soil-borne *Bacillus* spp. and *Pseudomonas* spp. against *Phytophthora infestans* and other potato pathogens. Front Microbiol 9:143.

85. Colombo EM, Pizzatti C, Kunova A, Gardana C, Saracchi M, Cortesi P, Pasquali M. 2019. Evaluation of in-vitro methods to select effective *streptomycetes* against toxigenic *fusaria*. PeerJ 7:e6905.

86. Zhang XD, Yang Q, Yu J, Zhao L, Fan JX. 2011. Research methods of single cell protein production from potato residues of *Bacillus pumilus*. Journal of Northeast Agricultural University 42:26–30.

87. Gao XY, Liu Y, Miao LL, Li EW, Hou TT, Liu ZP. 2017. Mechanism of anti-Vibrio activity of marine probiotic strain *Bacillus pumilus* H2, and characterization of the active substance. AMB Express 7:23.

88. Torres MJ, Brandan CP, Petroselli G, Erra-Balsells R, Audisio MC. 2016. Antagonistic effects of *Bacillus subtilis* subsp. *subtilis* and *B. amyloliquefaciens* against *Macrophomina phaseolina*: SEM study of fungal changes and UV-MALDI-TOF MS analysis of their bioactive compounds. Microbiol Res 182:31–9.

89. Zihalirwa Kulimushi P, Arguelles Arias A, Franzil L, Steels S, Ongena M. 2017. Stimulation of fengycin-type antifungal lipopeptides in *Bacillus amyloliquefaciens* in the presence of the maize fungal pathogen *Rhizomucor variabilis*. Front Microbiol 8:850.

90. Yu GY, Sinclair JB, Hartman GL, Bertagnolli BL. 2002. Production of iturin A by *Bacillus amyloliquefaciens* suppressing *Rhizoctonia solani*. Soil BiolBiochem 34:955–963.

91. Ongena M, Jacques P, Touré Y, Destain J, Jabrane A, Thonart P. 2005. Involvement of fengycin-type lipopeptides in the multifaceted biocontrol potential of *Bacillus subtilis*. Appl Microbiol Biotechnol 69:29–38.

92. Maget-Dana R, Ptak M, Peypoux F, Michel G. 1985. Pore-forming properties of iturin A, a lipopeptide antibiotic. Biochim Biophys Acta 815:405–9.

93. Yang MM, Wen SS, Mavrodi DV, Mavrodi OV, von Wettstein D, Thomashow LS, Guo JH, Weller DM. 2014. Biological control of wheat root diseases by the CLP-producing strain *Pseudomonas fluorescens* HC1-07. Phytopathology 104:248–56.

94. de Bruijn I, de Kock MJ, Yang M, de Waard P, van Beek TA, Raaijmakers JM. 2007. Genome-based discovery, structure prediction and functional analysis of cyclic lipopeptide antibiotics in *Pseudomonas* species. Mol Microbiol 63:417–28.

95. Ongena M, Jourdan E, Adam A, Paquot M, Brans A, Joris B, Arpigny JL, Thonart P. 2007. Surfactin and fengycin lipopeptides of *Bacillus subtilis* as elicitors of induced systemic resistance in plants. Environ Microbiol 9:1084–90.

96. Tran H, Ficke A, Asiimwe T, Hofte M, Raaijmakers JM. 2007. Role of the cyclic lipopeptide massetolide A in biological control of *Phytophthora infestans* and in colonization of tomato plants by *Pseudomonas fluorescens*. New Phytol 175:731–42.

97. Jourdan E, Henry G, Duby F, Dommes J, Barthélemy JP, Thonart P, Ongena M. 2009. Insights into the defense-related events occurring in plant cells following perception of surfactin-type lipopeptide from Bacillus subtilis. Mol Plant Microbe Interact 22:456–68.

98. Zhang X, Rui L, Lv B, Chen F, Cai L. 2019. Adiponectin relieves human adult cardiac myocytes injury induced by intermittent hypoxia. Med Sci Monit 25:786–793.

99. Li B, Li Q, Xu Z, Zhang N, Shen Q, Zhang R. 2014. Responses of beneficial *Bacillus amyloliquefaciens* SQR9 to different soilborne fungal pathogens through the alteration of antifungal compounds production. Front Microbiol 5:636.

100. Liu J, Hagberg I, Novitsky L, Hadj-Moussa H, Avis TJ. 2014. Interaction of antimicrobial cyclic lipopeptides from *Bacillus subtilis* influences their effect on spore germination and membrane permeability in fungal plant pathogens. Fungal Biol 18:855–61.

101. Tao Y, Bie XM, Lv FX, Zhao HZ, Lu ZX. 2011. Antifungal activity and mechanism of fengycin in the presence and absence of commercial surfactin against *Rhizopus stolonifer*. J Microbiol 49:146–50.

102. Grau A, Gómez Fernández JC, Peypoux F, Ortiz A. 1999. A study on the interactions of surfactin with phospholipid vesicles. Biochim Biophys Acta Biomembr 1418:307–19.

103. Cawoy H, Debois D, Franzil L, De Pauw E, Thonart P, Ongena M. 2015. Lipopeptides as main ingredients for inhibition of fungal phytopathogens by *Bacillus subtilis*/*amyloliquefaciens*. Microb Biotechnol 8:281–95.

